# Rapid Dopaminergic Signatures in Movement: Reach Vigor Reflects Reward Prediction Error and Learned Expectation

**DOI:** 10.1101/2025.03.24.645035

**Authors:** Colin C. Korbisch, Alaa A. Ahmed

## Abstract

Movement vigor across multiple modalities increases with reward, suggesting that the neural circuits that represent value influence the control of movement. Dopaminergic neuron (DAN) activity has been suggested as the potential mediator of this response. If DAN activity is the bridge between value and vigor, then vigor should track canonical mediators of DAN activity, namely learning signals in the form of reward expectation and reward prediction error. Here we ask if a similar time-locked response is present in vigor of reaching movements. We explore this link by leveraging the known phasic dopaminergic response to stochastic rewards, where activity is modulated by both reward expectation at cue and the reward prediction error at feedback. We used probabilistic rewards to create a reaching task rich in reward expectation, reward prediction error, and learning. In one experiment, target reward probabilities were explicitly stated, and in the other, were left unknown and to be learned by the participants. We included two stochastic rewards (probabilities 33% and 66%) and two deterministic ones (probabilities 100% and 0%). In both experiments, outgoing peak velocity increased with increasing reward expectation. Furthermore, we observed a short-latency response in the vigor of the ongoing movement, that tracked reward prediction error: either invigorating or enervating velocity consistent with the sign and magnitude of the error. Reaching kinematics also revealed the value-update process in a trial-to-trial fashion, similar to the effect of prediction error signals typical in dopamine-mediated striatal phasic activity. Lastly, reach vigor increased with reward history over trials, mirroring the motivational effects often linked to fluctuating dopamine levels. Taken together, our results highlight the link between known short-latency dopaminergic learning signals and the invigoration of movement, not only at the time of cue presentation and movement initiation, but during an ongoing movement immediately after feedback is provided.

**NEW & NOTEWORTHY:** Previous research has demonstrated the invigorating effects of reward on movement. Growing evidence suggests this is causally explained by midbrain dopamine transients. Here, we demonstrate that reach vigor tracks canonical variables of learning and motivation across time scales ranging from milliseconds to minutes. Velocity was modulated by reward expectation, reward prediction error and reward rate, key variables that have also been associated with striatal dopaminergic fluctuations. These results point to a potential neural mechanism by which dopamine can influence both decision making and movement control and support the proposition that reward-based invigoration of movement is in part influenced by dopaminergic circuits.

## INTRODUCTION

Imagine sitting down in front of a series of slot machines and being instructed to pull their levers, one after another. You find that these machines, frequently referred to as one-armed bandits for their ability to extricate money from patrons, have different rates of payouts, with some better than others. After learning these differences, you will likely prefer to pull the levers that offer a greater payout. But would you also choose to pull those levers faster or with more force?

Previous inquiry has shown that individuals will move faster towards goals or targets associated with greater value^1–10^. People are willing to produce greater muscle forces, or expend greater effort, to reach a cued target in less time. A potential explanation for how this may be represented in the brain lies in the neurotransmitter dopamine (DA), which is implicated in both the learning and representation of value as well as the control of movement^11^. Basal ganglia activity, already known to be influenced by dopaminergic inputs, has been found to invigorate not just movement vigor^12–15^, but the decision-making process as well^16,17^. If vigor is indeed a reflection of value and DA the mediator, then vigor may also reflect the machinery that underlies the learning process, due to the phasic dopamine release coincident with learning and reward prediction error^18^. Seminal work has shown that dopaminergic neuron (DAN) responses in a learned environment scale with reward prediction error at the time of both stimulus presentation and feedback presentation^19^. DAN activity is greater in response to cues associated with greater expectation of reward (greater positive reward prediction error), and lower upon feedback of that reward (lower positive reward prediction error). If a similar time-locked response is present in movement vigor at cue and feedback presentation, this would suggest that DA-related activity contributes to reward-driven increases in vigor.

While short-term phasic DAN response is implicated in prediction error and learning, tonic dopamine levels have been found to be sensitive to the history of reward reception^1,20–22^. Particularly, midbrain DA levels come to match average reward rate and are implicated in motivation and invigoration. Hamid et. al. found a significant relationship between history of reward and relative DA in the nucleus accumbens compared to other potential analytes. Greater tonic levels of DA can also be interpreted as greater motivation or *drive*, resulting in behavioral invigoration^23,24^. Recent reward history has been extensively recorded influencing movement vigor^20,25–28^. Returning to our hypothetical gambler, receiving more rewards over the past, regardless of which levers were pulled, would result in greater vigor in subsequent lever pulls.

A growing body of work has found correlates between phasic dopaminergic activity and movement vigor^29–37^. Cue-evoked phasic dopamine response within rat nucleus accumbens core (NAc core) was significantly correlated with trial movement vigor^30^. Similarly, for rat substantia nigra pars compacta (SNc) neurons sensitive to free movement initiation, relative activity levels were correlated with ensuing vigor^34^. Optogenetic activation of these neurons when not moving increased the probability of movement initiation, but did not produce any measurable changes in ongoing movement vigor. However, a separate study found that ongoing movement vigor was reduced with inhibition of striatal activity^33^. These and other works provide evidence that not only do longer timescale tonic dopamine levels potentially modulate animal movement vigor, so too may short latency, phasic responses.

In this study, we sought to investigate whether human kinematic response to probabilistic rewards would mirror the characteristic reward prediction error response observed in mesencephalic dopaminergic neurons (DAN) on a sub-second timescale. We also asked if kinematic response would reflect tonic DA response correlated with reward history. Given the short-latency and tuned response of DANs to reward value, we hypothesized vigor response in human arm reaching to be modulated by both the expectation of reward (probabilistic expectation) as well as the prediction error (difference between binary reward outcome and expectation). In effect, we expect outcomes congruent with two of the axioms for RPE-based models: 1) vigor responses to various reward expectations should demonstrate consistent lottery ordering, and 2) should show evidence of the “no surprise equivalence,” i.e., fully deterministic reward outcomes show no relative difference. We also hypothesized that greater reward amounts received over the recent past would lead to greater relative invigoration, as a result of increased striatal DA levels. To test these hypotheses, we performed two experiments, each of which involved human subjects performing out-and-back reaching movements to probabilistically rewarding targets. Our primary findings reveal that movement vigor is modulated online in alignment with reward prediction error on a sub-second time-scale, while also reflecting the learned value update on a trial-to-trial basis as well as average reward rate. These results highlight how motor planning, execution, and feedback control are all likely influenced by DA response, reflecting DA’s tripartite role in learning, motivation, and movement control.

## RESULTS

In a series of two experiments, we asked whether movement vigor tracked canonical determinants of dopamine-related learning and performance: learned value, reward prediction error, and reward history. In both experiments, human subjects directed arm reaching movements to targets associated with a probability of receiving a reward (Figure 1a-c; see Methods). Each of four target locations was associated with a unique probability of receiving the reward (0%, 33.3%, 66.6%, 100%) that was changed each block (Figure 1d) of 180 trials. Feedback of reward reception or reward denial was provided upon target acquisition. Assuming perfect representation of task instruction, participants could experience five reward prediction errors (RPE): −0.33 and +0.66 for the 66% target, −0.66 and 0.33 for the 33% target, and 0 RPE for the 0% and 100% targets (Figure 1e). This paradigm allowed us to probe the effect of reward expectation on the vigor of the outgoing movement. Additionally, we could investigate the effect of RPE at time of feedback on the vigor of the return movement. Lastly, we could test the effect of reward history on reach vigor across trials, independent of the expected value of the current target. We predicted that with increasing expectation, outgoing vigor would increase, reflected in an increase in peak velocity (Figure 1g), that vigor would be modulated on the return portion of the reach, scaled by the reward prediction error, and that vigor across trials would increase with reward history.

**Figure 1.**
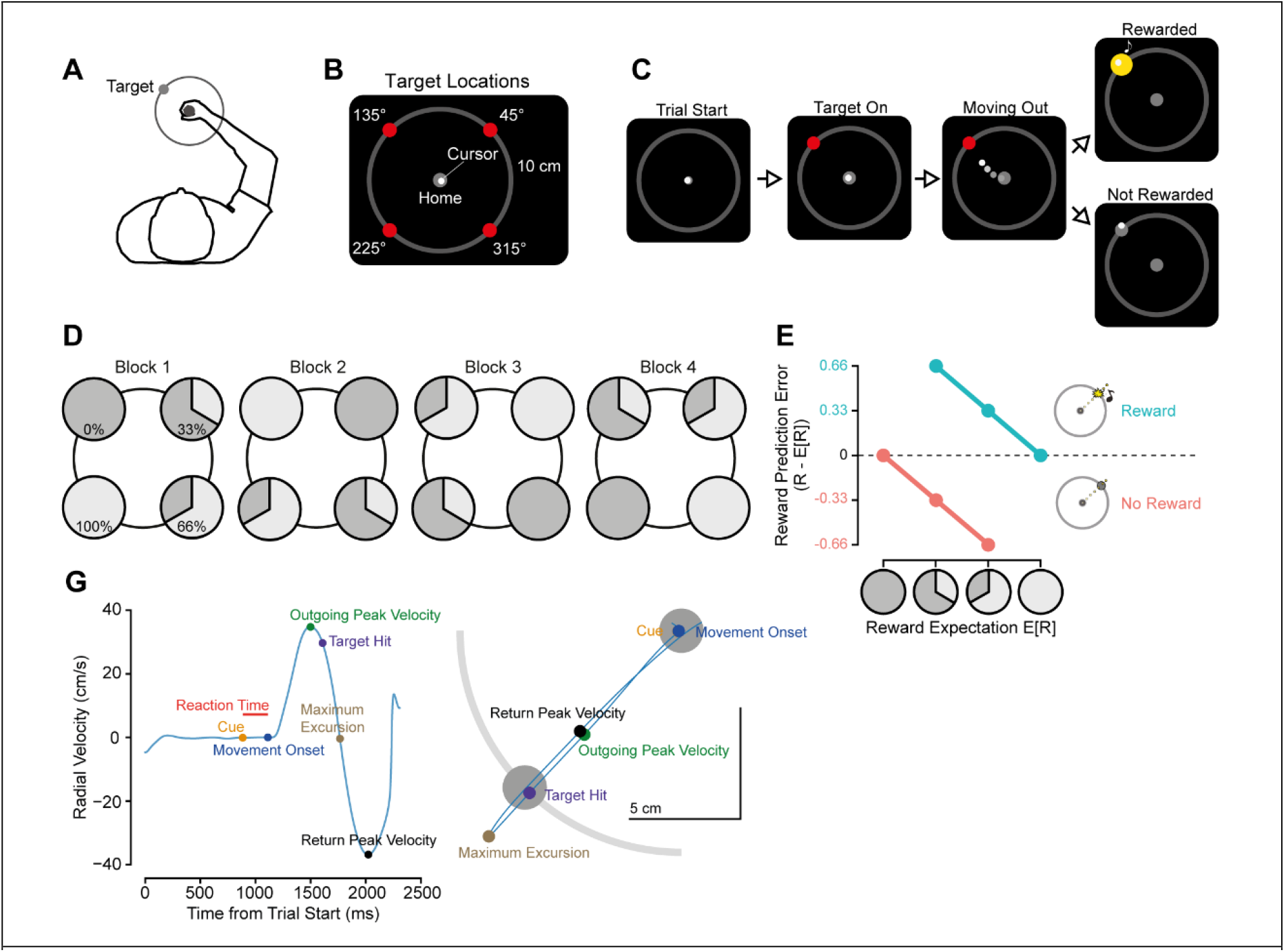
Experimental protocol. A) Participants made out-and-back reaching movements towards prompted, virtual targets. B) Targets were located at four different positions, each 10 cm from a central home location. C) Trial sequence. After the target was cued, participants were to make an out-and-back reach towards said target. Reward feedback was dependent on stated target reward frequency (in the first experiment), with reward feedback consisting of a yellow flash and high-pitched tone. Omission of feedback coincided with the target turning gray. D) At the beginning of each 180-trial block, in the first experiment, participants were told the reward frequencies of each target, either 0%, 33%, 66%, or 100%. Shown here is one possible experiment progression order. Relative order of reward-target associations was randomized across participants. Each block’s rewards were pseudorandomized such that, in the single-target trials, the target reward frequencies were exact to the underlying frequency. Each block consisted of 5 sets of 36 trials, wherein each target was presented 9 times, and amount of reward per target was precisely equal to the stated expectation (i.e., rewards were given 0, 3, 6, or 9 times within each set of 36 trials depending on the target). Target presentation order within each set of 36 trials was fully randomized. E) Reward prediction error (RPE) was defined as the difference between the stated (and actual) reward frequency per target and the binary outcome of the trial. G) *Left* Kinematic measures of interest included the time between target presentation (cue) and movement onset (reaction time), peak velocity of the outgoing portion of the movement, time between movement onset and target hit, peak velocity of the return portion, and difference between the absolute value of the two peak velocities. Example positional trace of trial reach trajectory given on *Right*, with spatiotemporal points of interest highlighted.

### Experiment 1

In the first experiment, we sought to focus solely on the vigor response to a known expectation of reward (E[R]). We did so by informing the participants of the reward probability of the targets at the beginning of each block.

### Peak velocity to target tracks reward expectation

As the cued target’s E[R] increased, peak velocity of the outgoing movement increased as well (GLMM, β^E[R]^=0.0159±0.00629, p=0.0152, Figure 2a,b). Likewise, time to target, defined as the time between movement onset and reaching a 10 cm radial distance, decreased with increasing reward expectation (β_E[R]_=−0.0125±0.00527, p=0.0226). Reaction times decreased with greater E[R] (Supplemental Figure 1a-b).

**Figure 2.**
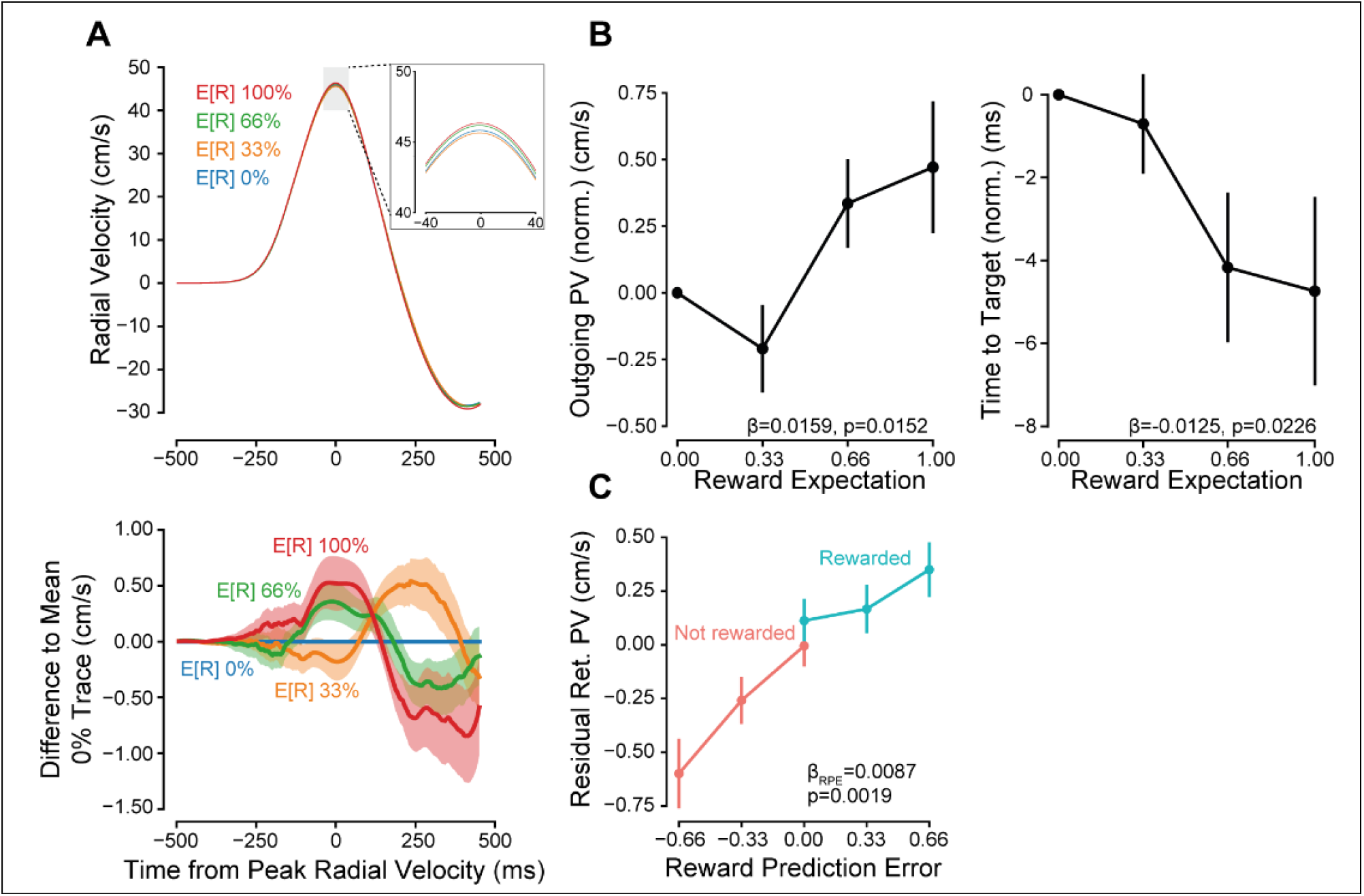
Vigor tracks reward expectation. A) *Upper* Radial velocity trace for first experiment, aligned to time of outgoing peak velocity. Highlighted region shows differences in peak excursion velocity for differences in reward expectations. *Lower* To calculate the difference traces, each participant’s average signed radial velocity, at each sampled time point for the 0% expected reward condition, was subtracted from all other trials. Traces shown are the grand average of these difference traces, +/-standard error. B) *Left* Outgoing peak velocity increased with increasing expected reward. Data were normalized by subtracting the per-participant average for 0% expected reward condition. *Right* Similarly, the time to target hit, defined as the time between movement onset and reaching out to a 10cm radial distance, decreased as reward expectation increased. C) Return peak velocities were residualized by first fitting a reduced model including target direction and outgoing peak velocity. These residuals were significantly correlated with trial reward prediction error, increasing with greater RPE.

### Return velocity is influenced by reward prediction error

When the cursor hits the target, participants can either receive reward feedback or not. Does this feedback, or lack thereof, influence subsequent movement kinematics? Does the degree of surprise also modulate movement kinematics? In other words, does the reward prediction error influence the speed of the ensuing movement?

To answer these questions, we analyzed participants’ return movements and whether receiving (or not receiving) reward feedback on a given trial produced significant influences. Specifically, we questioned whether the sign and magnitude of the reward prediction error, *RPE*, influenced the velocity:

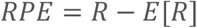

with *R* a binary coding for presence or absence of reward feedback (0,1), and *E*[*R*] the instructed expectation of reward feedback (see Methods).

Participants exhibited diminished peak velocity on return movements compared to the outgoing movements (paired mean difference = 0.0601 m/s [0.0445,0.0757 95% CI], t_41_=7.7789, p=1.34e-9). After controlling for the average relationship between outward and return peak velocities (β_OutPV_=0.580±0.0192, p<2e-16), we observed an effect of reward feedback and reward expectation, i.e., RPE (β_RPE_=0.00865±0.00260, p=0.00186). This indicated that the return portion of the reach was not merely driven by feedforward mechanisms, but also feedback. No significant main effect of reward was found (β_Reward_=0.000876±0.00159, p=0.586), nor interaction with RPE (β_RPE×Reward_=−0.00545±0.00392, p=0.171), signifying a slope of response across prediction error values without discontinuities. Additional control analyses confirmed return movements were modulated by reward feedback and not other potential confounds (Supplementary Figures 2-4).

To better account for variance in the outgoing portion of the movement, we normalized instantaneous velocity and focused on within-trial effects. First, instantaneous radial velocity was divided by the within-trial outgoing peak velocity. Next, the difference traces of these %-outgoing velocity values were calculated in a similar manner as in figure 2A, though instead of taking the difference to the 0% reward trials, the differences were taken relative to the per-participant averaged normalized velocity trace for all 0% and 100% reward trials, i.e. RPE=0 trials. After calculating the grand average of these difference traces, we see a clear striation in relative normalized return velocity corresponding with reward prediction error (Figure 3a).

**Figure 3.**
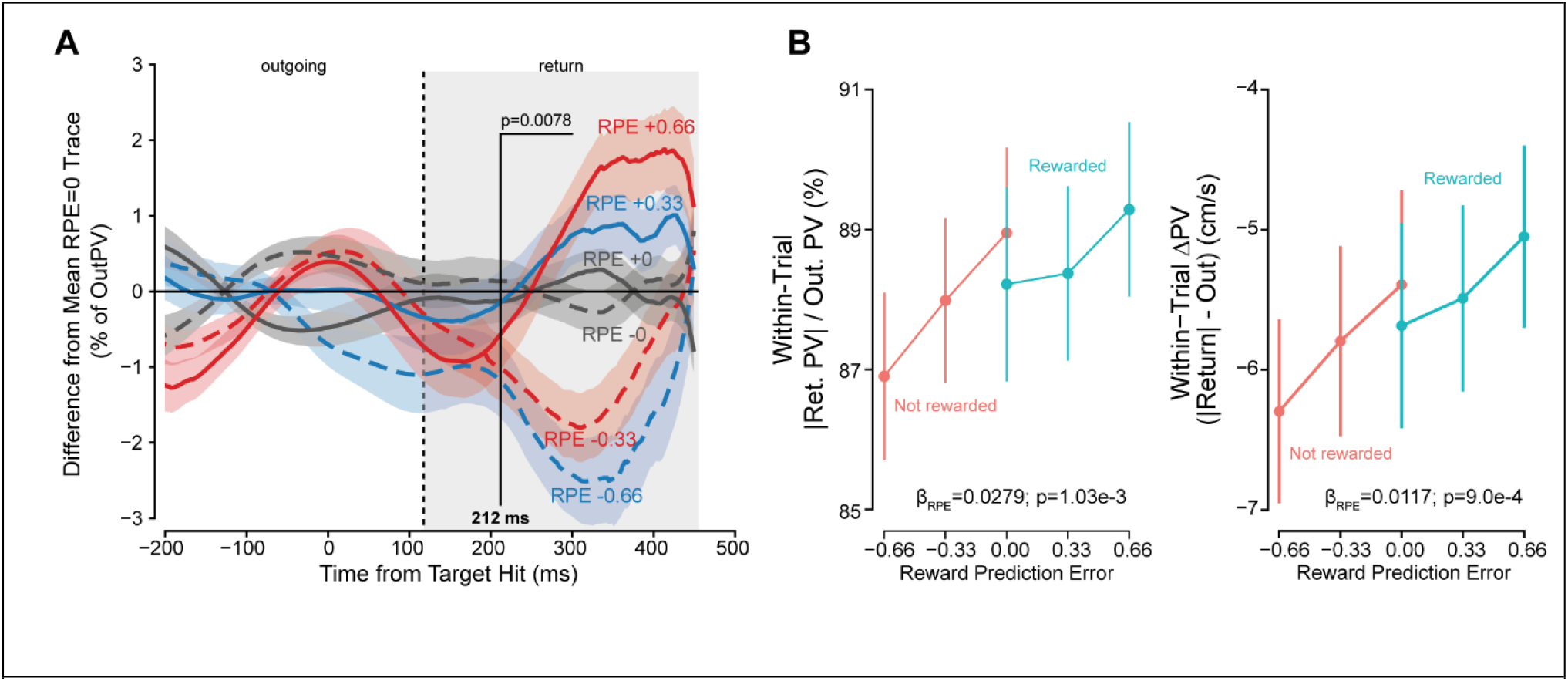
Vigor response reflects reward prediction error. A) For a within-trial measure of relative invigoration, instantaneous velocity was normalized by dividing by the trial outgoing peak. Difference traces were then calculated in a manner similar to figure 2A. From SPM analysis, a significant effect of RPE on percent peak velocity was found beginning at 212 ms after reward feedback was received. B) The within-trial difference in absolute peak velocities, *Left* return divided by outgoing *Right* return minus outgoing, significantly varied with reward prediction error. As RPE increased, the return peak more closely matched the outgoing peak.

We next performed a hierarchical random effects analysis (Supplementary Figure 5; see Methods) to determine the population-average effect of reward prediction error on instantaneous, normalized velocity. At 212 ms after target feedback, we found a significant negative effect of RPE, indicative of greater invigoration in the return movement with greater RPE (Figure 3a).

Aside from the effect on instantaneous velocity, the within-trial difference in velocities, i.e., the difference between excursion and return peak velocity, also exhibited a significant effect of RPE (Figure 3b); with the slope of response consistent across positive and negative RPE conditions (β_RPE x Reward_ =−0.00176±0.00502, p=0.727). For both percent relative to outgoing peak (3b *Left*) and velocity difference (3b *Right*) the slope of response was significant across RPE conditions (β_RPE_=0.02788±0.00788, p=1.03e-3; β_RPE_=0.0117±0.00325, p=9.00e-4, respectively). When RPE was most positive (+0.66) the difference between excursion and return peak velocity was at its smallest compared to RPE at its most negative (−0.66). There was no separate effect of reward feedback in either metric (β_Reward_=−0.00269±0.00463, p=0.5643; β_Reward_=−0.00355±0.00208, p=0.0978).

Overall, we found a significant effect of reward feedback on the return movement that was dependent on the reward expectation, i.e., the reward prediction error. With greater RPE, relative return velocity was greater compared to when the prediction error was negative. This difference emerged 212 ms after feedback was received and could not be accounted for by variation in maximum excursion, nor preemptive differences in the outgoing portion of the movement (potential correlations between outgoing peak velocity and future reward reception). Participants employed neither set return velocity, nor set return velocity difference strategies. In the first case, outgoing peak velocity would be unrelated to the return peak, and in the second case, return velocity difference would be invariant to the experienced prediction error.

### Experiment 2

We next asked whether vigor was modulated by learned value, rather than instructed value. To probe this we conducted a second experiment where participants were left uninformed of the targets’ reward frequencies. Instead, participants learned through experience over the course of a block, affording us a measure of trial-to-trial response of changes in reward estimation. Within each block, after a sequence of 144 single-target trials similar to the first experiment, there were 36 two-alternative forced choice trials (2-AFC) to assess degree of learning (Figure 4a). Individuals were incentivized to choose greater expected rewards on choice trials as additional monetary compensation was dependent on performance (see Methods). Critically, no feedback was given on target hit during these choice trials to limit learning to the single target trials only.

**Figure 4.**
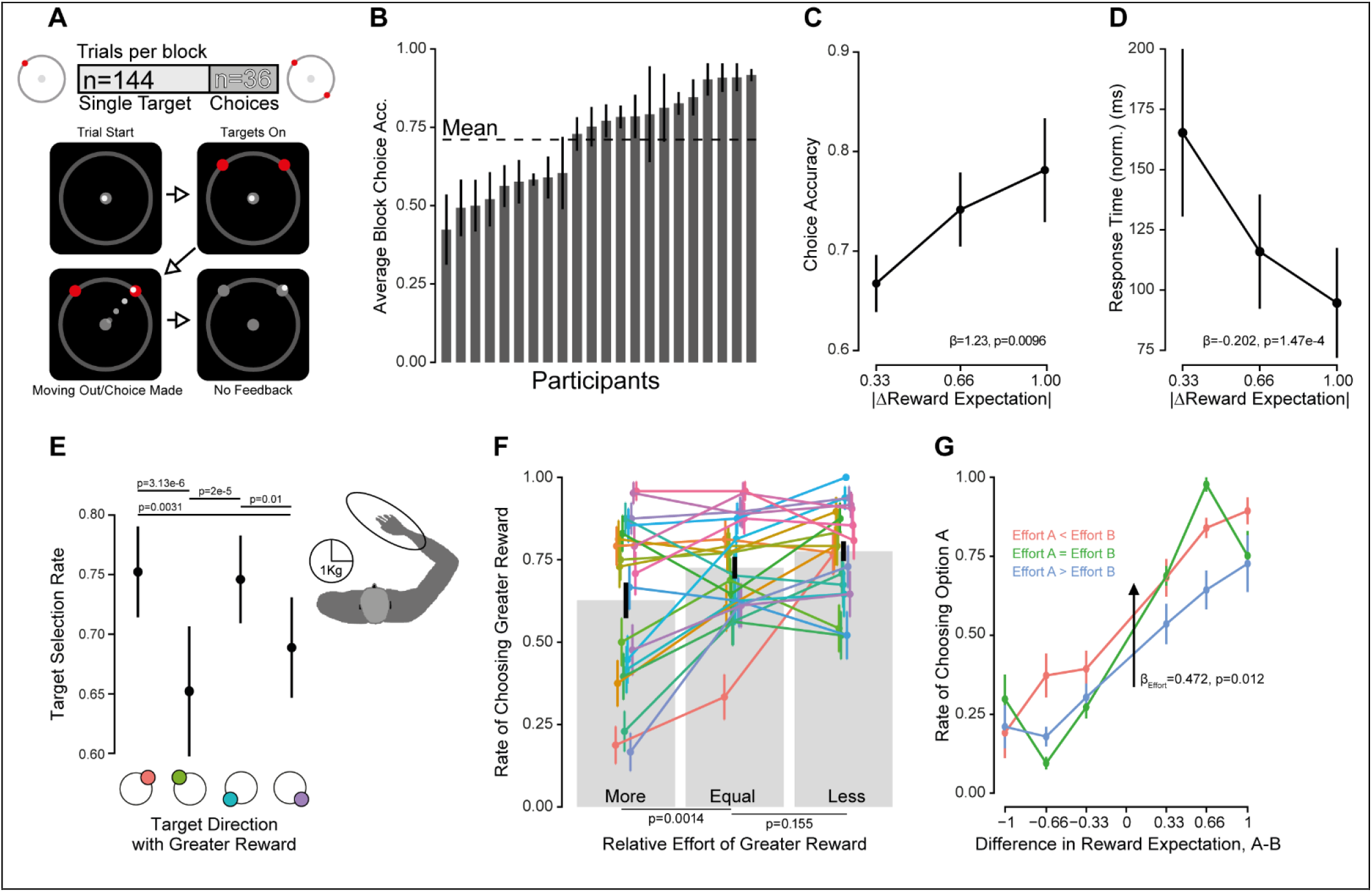
Experiment 2 choice behavior. A) In the second experiment, reward probabilities were left unstated to the participants. Instead, they were incentivized to learn the underlying rewards during the single-target period, as additional monetary compensation was dependent on performance during later choice trials (see Methods). B) Per-participant average accuracy rate over four blocks. Participant choices were highly variable across the population. Mean accuracy was 71%. C) Participant choices were found to be sensitive to relative expectation, choosing the more rewarding option as difference increased. D) Response time decreased with increasing difference of reward expectations of the presented targets. E) Target direction was effected choice selection. When targets 1 or 3 were the more rewarding option, i.e. associated with lesser biomechanical effort, they were selected at higher rates compared to when targets 2 or 4 were the more rewarding option, consistent with a potential influence of biomechanical effort on choice selection. F) When grouping choices by whether the greater reward has greater relative effective mass, significant differences in choice accuracies were found. Individual participant responses are shown in color. G) Baseline selection rates were shifted depending on relative effort. No interaction between effort and reward difference was found.

### Choices reflected both expectation of reward and effort

Choice accuracy, defined as the selection of the greater reward frequency of the two options, varied across the participants, ranging from 42.7% to 91.7% (Figure 4b). The population-average accuracy rate was 71%. Underlying reward difference, hidden to participants, was found to significantly predict rate of choosing one option over another (β_ΔE[R]_=1.233±0.476, p=0.0096; Figure 4c). Response time of the decision was found to significantly decrease with increasing difference of reward expectation (Figure 4d; β_|ΔE[R]|_=−0.202±0.053, p=1.47e-4), providing further evidence that underlying expectations were learned. Interestingly, response time and outgoing peak velocity response differed during choice trials in their sensitivity to reward difference and choice accuracy (Supplementary Figure 6). Whereas outgoing peak velocity only varied with the selected option’s reward expectation, response time varied with both relative and selected expectation.

Target direction was also found to influence decision making. We first categorized all choices by whether a given target direction was associated with the greater of the two potential rewards presented on a given choice trial. We then calculated the choice accuracy rate for each of these subsets. If direction had no influence, we should expect no variation in selection rates between these four categories, however this was not the case. Instead, significant differences were found (Figure 4e). Selection rate biases matched the arm’s inertial axes^38,39^ (Figure 4e *Right*). Using this measure of effort, we categorized choice trials by whether the more rewarding option was associated with more or less effort compared to the alternative choice (Figure 4f). We found significant differences in accuracy rates, with idiosyncratic responses to effort readily apparent. Augmenting our logistic regression analysis from before by including a relative effort term, we found a significant effect of said relative effort, with average rates of choosing the more rewarding option increasing when this option was associated with less effort (β_e_=0.472±0.188, p=0.012; Figure 4g). Thus, participants’ choices revealed they had largely learned the target reward contingencies and demonstrated that these value-based choices were influenced by the effort of the arm reach.

### Velocity in single-target trials tracked reward expectation and reflected learning

Given that participants’ choices revealed a learned reward estimation, we asked whether vigor on single target trials would reveal this estimation as learning progressed. Over the course of the single-target period, we found that average slope of response of outgoing peak velocity relative to reward expectation increased (GLMM; β_Trial x E[R]_ = 0.0392±0.01447, p=0.00682), demonstrating a dynamic response to the probabilistic reward (Figure 5a). Likewise, the effect of expectation on the time to target significantly varied over this period (β_Trial x E[R]_ = −0.0356±0.0142, p=0.0125). At the beginning of a block, there was no significant effect of reward expectation (β_E[R]_= −0.0108±0.0073, p=0.142). The change in response of instantaneous velocity over the course of the single-target period is further evident in the velocity difference traces (Supplementary Figure 7). However, no such interaction effect between trial and hidden reward expectation was found when modeling reaction times (Supplementary Figure 1).

**Figure 5.**
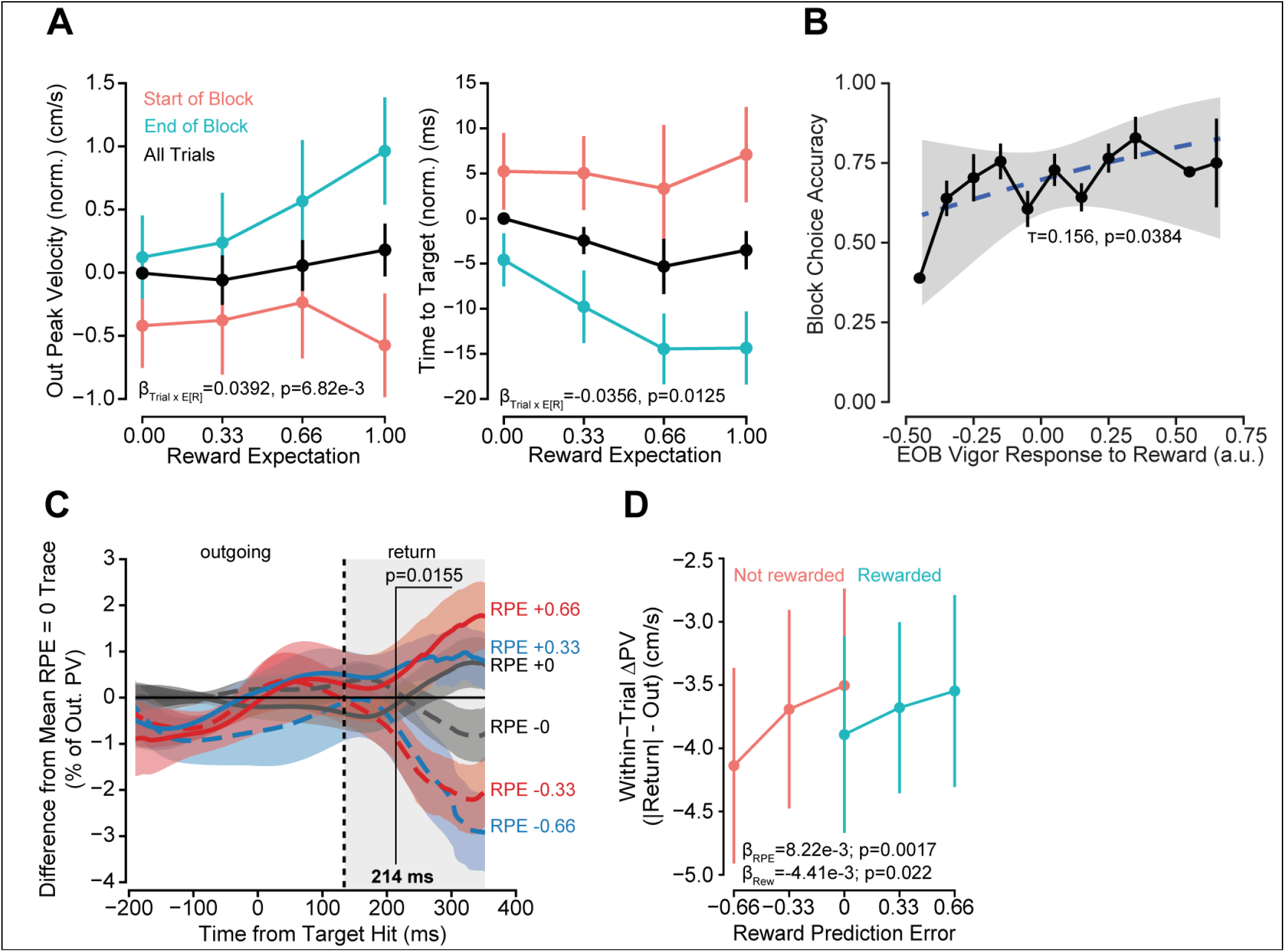
Experiment 2: Vigor tracked reward expectation and reward prediction error. A) *Left* Outgoing peak velocity increased with increasing reward expectation. Data were normalized by taking the difference to per-participant average for 0% reward. Points and error bars represent grand means, ± standard error. The slope of peak velocity response to reward expectation changed over the course of a block. In the initial 36 trials (start of block), response was essentially flat, but by the final 36 trials (end of block), outgoing velocity significantly increased with respect to reward frequency. *Right* Time to target, taken as the duration between movement onset and reaching out to a 10 cm radius (target hit) changed with respect to underlying expectation over the course of the single-target period. B) End of block response to reward (i.e. per-participant slope of response to E[R]) was significantly correlated with block choice accuracy (rate of choosing the more rewarding option, Figure 4B). C) Normalized velocity difference traces for experiment 2. As in figure 3C, velocity data were first normalized by dividing by the within-trial peak outgoing velocity. Difference traces were calculated by subtracting from each trial the average instantaneous normalized velocity for trials where the underlying reward expectation was 0% or 100% (RPE = 0). Data presented is the grand average of these difference traces. Differences in relative velocity emerged after reward feedback that scaled with prediction error. D) As RPE increased, the relative reduction in peak velocities between the return and outgoing portions of the reaching movement decreased (i.e. the difference was less negative).

The strength of peak velocity-reward expectation response (i.e. the slope of velocity relative to reward) could predict within-block choice accuracy (the rate of choosing the more rewarding of the two options). Across the participant pool, we found a significant correlation between these measures (Kendall’s rank correlation, *τ* = 0.156, p=0.0384; Figure 5b). The stronger the slope of response in an individual at the end of the single target period, the greater the rate in selecting the more rewarding of the two options presented during choice trials.

Even when participants were not told of the expected reward frequencies at the outset of a block, both outgoing peak velocity and time to target responded to increasing expectation. Learning of the different reward frequencies is shown clearly by the differential response of velocity at the beginning and end of the single-target period. Initially, kinematic response did not differ between the four targets, but by the end, average peak velocity increased with increasing expectation of reward feedback. Within-block performance could be predicted based on participants’ slope of response to reward at the end of the single-target period, suggesting that the change in kinematics over the course of a block was related to the learning of relative reward frequencies.

### Change in return velocity tracks reward prediction error

Turning to within-trial measures of relative vigor, we performed the same hierarchical random effect SPM analysis on normalized velocity as in the first experiment and found a population-level effect of RPE (Figure 5c). Additional supporting detail for this analysis is provided in supplementary documentation (Supplementary Figure 8). At 214 ms after reward feedback, relative velocity varied significantly with reward prediction error. Likewise, the within-trial velocity difference (return minus outgoing) also significantly varied with RPE (β_RPE_=8.22e-3±2.61e-3, p=0.00166; Figure 5d). With increasing positive prediction error, the return peak velocity more closely matched the outgoing peak velocity. Reward feedback had no significant interaction effect on this difference (β_RPE x Reward_ =−2.156e-3±3.682e-3, p=0.558), However, there was a significant main effect of reward (β_Reward_ =−4.414e-3±1.825e-3, p=0.0217). As such, for both 0% and 100% rewarding trials, the differences in outgoing and return peak velocities were statistically distinct. The slope of response to reward prediction error was not found to change over the course of a block (β_RPE x Trial_ =−4.746e-3±4.770e-3, p=0.320). As in the previous experiment, we performed additional control analysis to isolate the effect of reward feedback on the return movement (Supplementary Figure 9).

To reiterate, even when expected reward must be experienced and learned, individuals change their behavior on the return movement in a manner consistent with RPE sign and magnitude.

### Biomechanical effort slowed outgoing peak velocity

Considering the effect of target direction (i.e. relative effort), on choice bias, we were also concerned whether there would be differences in average peak velocities towards the different directions. Investigating this potential effect (Figure 6a *Left*), we found that each direction was significantly different in average peak outgoing velocity from the other (Tukey-HSD comparison testing; Supplementary Table 2). In addition to the directional effect, we found that average outgoing peak velocity increased over the course of the experiment from block-to-block (β_Block_=0.158±0.051, p=0.0054).

**Figure 6.**
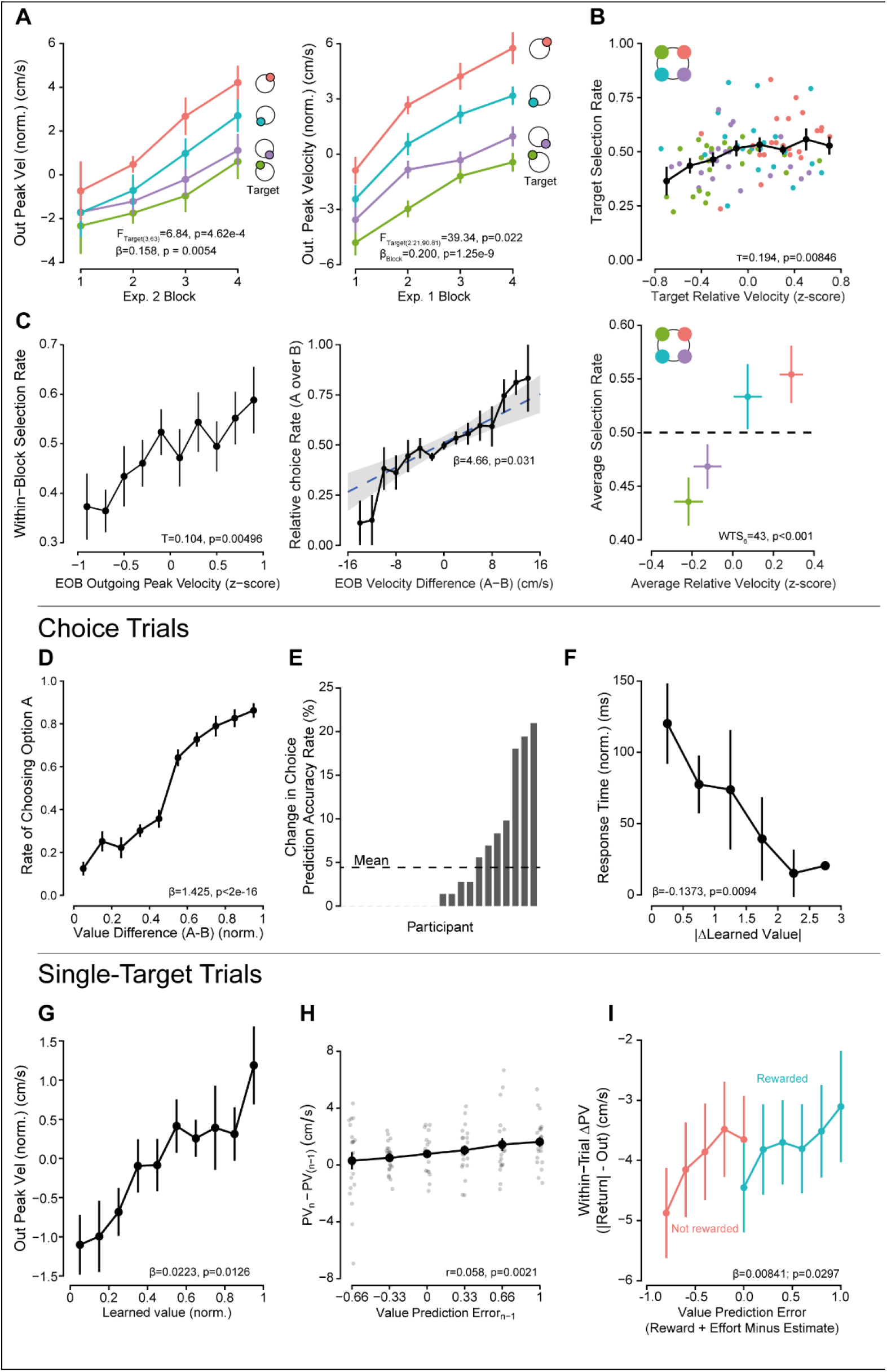
(Top) Relation between velocity, subjective effort, and learned value. A) *Left* In experiment 2, arm reaches towards target 1 were found to have a greater peak velocity on average, with targets 2 and 4 associated with comparatively lesser averages. *Right* In the first experiment, the same ordering of relative peak velocities was observed. B) *Upper* Participant-relative velocity, in single-target trials, were correlated with relative rates of selection of the different target directions. *Lower* Averaged over all participants per target, there was evidence of a separation in relative average velocities and selection rates. C) *Left* Relative velocity to a given target at the end of the single-target period could predict preference. *Right* Difference in average velocity between two targets, within the final 36 single-target trials, was continuous with relative rates of selection for those two targets. The greater the difference in velocity, the more frequently one option was chosen over the other in the proceeding six choice trials where the two targets were presented together. (Bottom) Performance of Learned Value Model D) Individuals were sensitive to learned value differences during decisions, selecting targets with greater relative values. E) Choice prediction accuracy improvement was participant specific. Average improvement was 4.42%, and some participants saw choice prediction improvements up to 20% (roughly 29 choices during the experiment). F) Choice trial response time significantly decreased with difference of the presented targets’ learned values. G) Normalizing value relative to per-participant minimum and maximum estimate (min-max norm.), we found that outgoing peak velocity during the single-target trials significantly increased with increasing learned value. H) On trials where the same target was prompted in repetition, the previous trial’s reward prediction error was significantly correlated with the trial-to-trial change in outgoing peak velocity. I) Value prediction error, taken as the difference between feedback and expected value, was also found to predict the relative difference in peak velocity between outgoing and return.

We also evaluated whether this directional influence on velocity was conserved in the previous experiment (Figure 6a *Right*). Once again, differences in average outgoing velocity across the directions were consistent with the effective mass of the arm. Peak velocity increased over the course of the experiment (β_Block_=0.2352±0.0058, p=1.25e-9) for all targets, and post-hoc multiple comparison testing (Tukey HSD) revealed significant differences across all four targets (Supplementary Table 1).

In short, average velocity to the different target directions in both experiments reflected the relative inertia, or effective mass, of the arm when making this reaching movement. Outgoing peak velocity was greatest towards the 45° target, and slowest towards the 135° target.

### Subjective vigor response in single-target trials predicted choice

Summarizing our results thus far for the second experiment, we have shown how participant choices reflected subjective value, and kinematic response during single-target trials reflected the learning of relative reward frequencies. To further substantiate the relationship between vigor and preference, we examined the degree to which reach velocity during the single-target trial period could predict choices across our participant pool.

To start, we analyzed whether the subjective effort bias evidenced in participant choices would also be present in the kinematic response. Comparing the per-participant relative velocities and the target selection rates, we found a significant correlation (Kendall’s rank correlation, *τ* = 0.194, p=0.00846; Figure 6b *Left*). With MANOVA analysis, after averaging across participants, we found significant differences between the relative velocities and selection rates across the four target directions (Figure 6b *Right;* WTS_(6)_=43.165, p<0.001). Next, we questioned whether kinematic behavior during the end of the single target period (the final 36 single-target trials) could predict later choice selections directly. To do so, we analyzed the correlation between the relative peak velocity per-target, per-participant and that target’s within-block selection rate (quantified as a fraction of the total 18 potential selections within the choice period). A significant correlation was found (Kendall’s rank correlation, *τ* = 0.104, p=0.00496; Figure 6c *Left*). The faster someone reached towards a target at the end of the single-target period, the more frequently they chose that option during choice trials.

As an additional piece of evidence for the relationship between outgoing peak velocity and relative selection rate, we directly compared the difference in mean velocity between two targets at the end of the single target period and the relative rate of selection within that block. Via logistic regression analysis, we found a significant relationship between the velocity difference and rate of selection (β_ΔPV_ = 9.56±2.55, p=1.75e-4; Figure 6c *Right*). Roughly, a 0.2 m/s difference in peak reach velocity translates to a 3:1 preferred rate of selection.

Taken together, we found that participant decisions varied widely across the population but could be predicted by each participant’s relative outgoing velocities on single target trials: the greater the relative peak velocity, the more frequently that option was chosen.

### Value Estimation

Reward and effort (in the form of target direction) had significant influences on outgoing velocity. Both reward and effort also affected choice preference and relative selection rates. We posit that the target characteristics of reward expectation and direction could be combined into a singular subjective value or decision variable that would describe both participant-specific reach velocity and choice preference.

### Learned value better explained single-target trial reach vigor

To examine whether this was the case and estimate this subjective decision variable, we fit a Bayesian hierarchical delta-rule learning model, similar to a Rescorla-Wagner model_40_, to choices at the end of each block to determine the per-trial subjective value for each participant. After each trial, the updated value estimate of a target *V′^T^* is calculated:

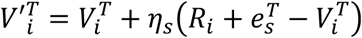

Here, *R_i_*is the reward received on the current trial, *η_s_*is learning rate, and 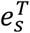, the relative effort of the target, dependent on direction (major or minor inertial axis). Participant-specific parameters, *η_s_*and 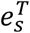, were selected and sampled from an estimated population distribution in a hierarchical fashion.

A separate logistic regression confirmed that value differences predicted choice behavior (Figure 6d). As relative value increased, individuals were more likely to choose that option. Additionally, the model-derived learned value was found to better predict choices as compared to reward expectation, with an aggregate accuracy rate of 75.4% compared to 70.98%. This was corroborated with a significant difference in ROC curves between the two metrics (Bootstrap test for difference in AUC: D=8.054, n=2000, p=8.014e-16). However, prediction improvements were highly idiosyncratic in the participant pool (Figure 6e). Lastly, choice trial response time was significantly influenced by value, decreasing with increasing value difference (β_|ΔV|_=−0.1373±0.0464, p=0.0094; Figure 6f). Outgoing reach during choice trials was also influenced by the value estimate (Supplementary Figure 10).

We next used this learned value estimate to predict reach peak velocity on single target trials. There was found to be a significant relationship between value and outgoing peak velocity (GLMM; β_Value_=0.0223±0.00895, p=0.0126; Figure 6g). The regression model including value improved in performance relative to one using E[R] (AICc: −13479.4 versus −13369.3 respectively). Likewise, time to target decreased with learned value as well, and better matched the data compared to reward expectation (β_Value_=−0.0265±0.007148, p=0.0013; Figure S6B; AICc: −12225.6 versus −12157.7).

Interestingly, even though subjective directional efforts were incorporated into the target value, there were still significant effects of direction. Post-hoc Tukey HSD testing revealed significant differences between the target-specific regression factors (Supplementary Table 3). This implies that the difference in vigor between directions does not fully capture the resultant difference in value as shown by later choices.

We predicted, as an additional piece of evidence relating learned value and vigor, that an individual’s change in value (the value prediction error) would be correlated with the trial-to-trial change in peak velocity after a target is repeated (Figure 6h). Calculating this correlation, we found that, as hypothesized, a greater value update leads to greater velocity changes, and the relationship was significant (Pearson’s correlation, t_(2801)_=3.083, r=0.058, p=2.07e-3). Testing to ensure this effect was target-specific, we calculated this correlation on trials where relative effort, but not direction, was repeated, and found no significance (t_(2732)_=−0.624, r=−0.012, p=0.533).

The relative difference in peak velocities between the outgoing and return portions of the movement decreased with increasing value prediction error (VPE) (β_VPE_=0.008414±0.00370, p=0.0297; Figure 6i); the comparative velocity of the return portion of the reach was greater with more positive error. Performance in modeling the within-trial velocity difference also improved when using VPE compared to RPE (AICc: −37118 versus −37098.2). Unlike the RPE regression model that showed a significant effect of reward reception on average difference in return velocity (see above, Figure 5d), the value prediction error model did not (β_Reward_=−0.00441±0.00296, p=0.153). Additionally, no significant interaction was found (β_VPE x Reward_ = −0.00510±0.00298, p=0.0873). In effect, the slope of response of velocity difference to value prediction error remained constant with or without reward feedback. Taken together, relative return velocities varied continuously with value prediction error with no additional significant effect of reward reception.

### Recent history of reward led to faster movements

Having shown how target-specific learned value influenced the reach kinematics on the outgoing and return portions of the movement, we sought to investigate the potential effects of target-agnostic reward reception, independent of value estimation. We investigated the potential effect in both experiments, testing the influence of previous trial reward reception and an integrated reward history, making use of the “leaky integrator” model^41^ (see Methods). Based on previous findings, we hypothesized that reward history would invigorate the outgoing movement, increasing peak velocity, and decrease response times.

In our first experiment, we found an effect of previous reward reception on outgoing peak velocity. Peak velocity was greater on trials following reward compared to trials following no reward (β_PriorR_=0.0152±0.003, p=5.93e-6; Figure 7a). For our reward history model, after fitting to reaction times, the smoothing factor, *α*, was found to equal 0.647. Incorporating reward history into our previous regression model for outgoing peak velocity, we found that velocity significantly increased with increasing 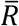 (β_History_=0.0209±0.00319, p=7.15e-8; Figure 7b). Model performance also relatively improved (AICc: −85753.7 versus −85728.6). This model-derived reward history value was significant even when accounting for current trial reward expectation (Figure 7c, Supplementary Figure 11).

In our second experiment, unlike the first, there was no significant response caused by reward reception on the previous trial (β_PriorR_=−0.0033±0.006, p=0.592; Figure 7d). With the same reaction time dependent likelihood maximization, we found a smoothing factor, *α*, equal to 0.0521. Outgoing peak velocity again significantly increased with reward history (β_Hist_=0.0884±0.0151, p=4.66e-9; Figure 7e, Supplementary Figure 11e). Model performance when predicting velocity was also improved (AICc: E[R]+History −36774.7, E[R] −36742.4).

**Figure 7.**
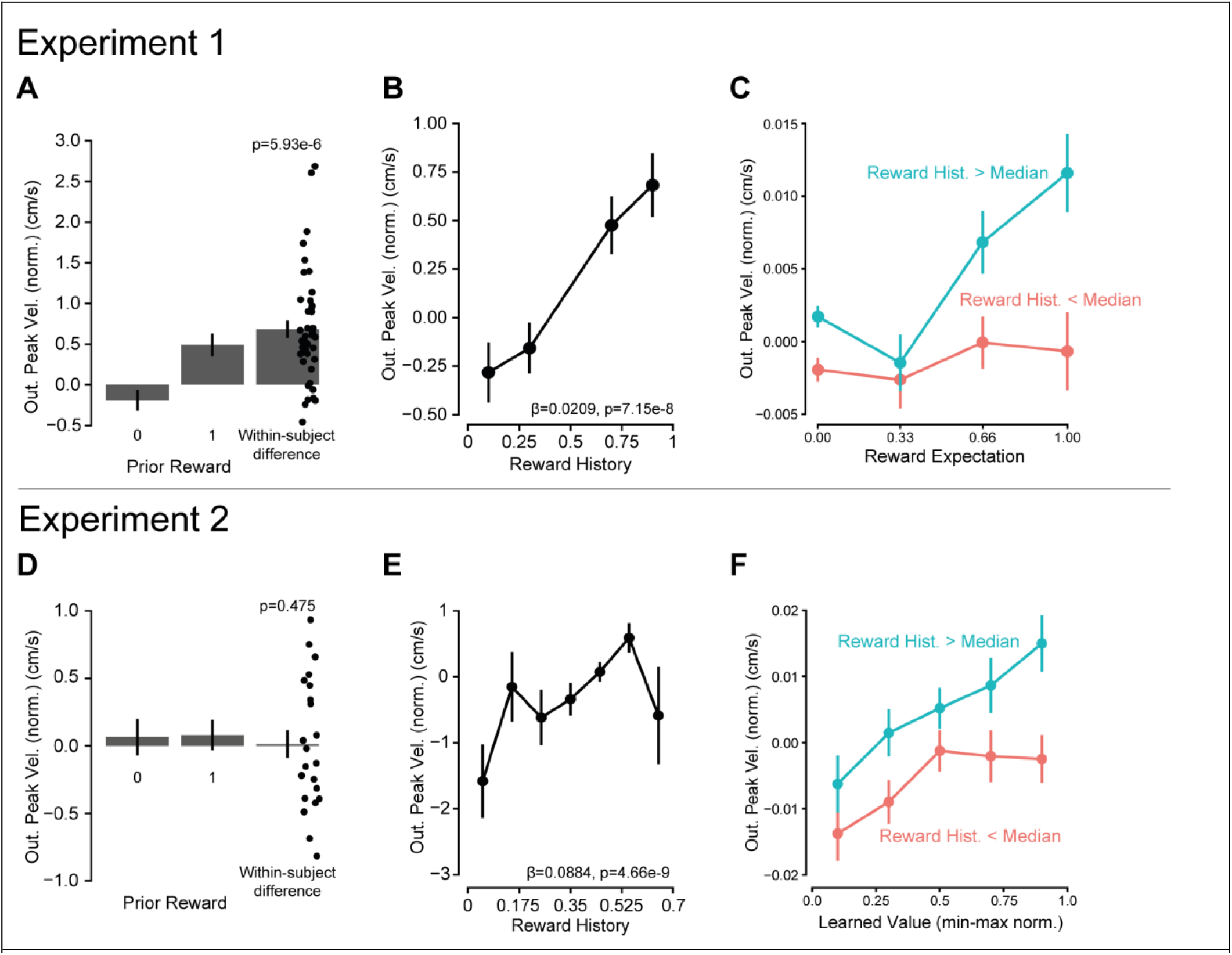
Peak velocity increased with reward history. Experiment 1: A) Outgoing velocity significantly increased with prior reward reception B) Reward history was estimated based on an exponentially smoothed moving average fit to reaction time (see Methods). With greater reward history, outgoing peak velocity increased. C) Even when accounting for movements to the same reward expectation, reward history has a significant effect on outgoing peak velocity. Experiment 2: D) Prior reward reception did not influence the subsequent trial velocity. E) The effect of reward history was determined in the same manner as in experiment 1. Outgoing peak velocity significantly increased with the amount of reward received in recent trial history. F) Vigor of reaching movements towards targets of similar learned value, were influenced by recent trial reward history.

Lastly, including both learned value and reward history in a single generalized regression model significantly improved model likelihood, compared to an alternative model that excluded the reward history term (LRT: 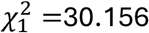, p=3.987e-8). This result strengthens our argument that both recent, target-agnostic rewards and effort-conscious valuation of specific targets influence the motivation of movement.

## DISCUSSION

Here, we demonstrated that reach vigor tracks neural correlates of learning and motivation across time scales ranging from milliseconds to minutes. Velocity was modulated by reward expectation, reward prediction error and reward rate, key variables that have also been associated with striatal dopaminergic fluctuations. These results point to a potential neural mechanism by which dopamine can provide the bridge between decision making and movement control.

Reward prediction error is typically examined with respect to recent reward history and current target expectation. Saccadic reaction times are faster or slower following a positive or negative reward prediction error^42^. In this study, we are the first to show RPE can have a modulatory effect of an ongoing movement, leading to either acceleration or deceleration that aligns with the sign of the prediction error. Moreover, we find that this effect scales with the magnitude of the prediction error. The rapid reflection of RPEs in vigor may be related to vigor’s already known response to average reward rate of the environment^27,43^, which is tracked by tonic dopamine levels^21,26^. Striatal dopamine levels rapidly rise and fall as a consequence of the temporal integration of phasic prediction error signals^32,44^. It could be through this mechanism that we see rapid sub-second changes in vigor. However, the exact relationship between spiking activity and striatal dopamine levels is multifaceted^29,45^ and the impact we observe on vigor is small and fleeting. This may explain why targeted optogenetic manipulations were limited or temporally restricted in their effects on vigor^34^.

A substantial body of evidence has detailed the relationship between midbrain DA transients and action initiation^46^. In both freely-moving, and head-fixed mice, phasic activity accompanied movement initiation^34,47^, with its correlation to the ensuing movement’s vigor being context dependent. Likewise, many studies found significant relationships between phasic DAN activity and reward-seeking motor behaviors^20,35,48^.

It may seem unlikely that the absence or presence of the reward feedback visual stimulus would influence ongoing velocity at sub-second speeds, approximately 215 ms after stimuli presentation. However, neurons in the superior colliculus, that then project to dopaminergic neurons within the VTA or SNc, have a response latency between 40 and 60 ms^49^. Another study found median response latencies of cue-elicited DAN responses within the SNc were on the order of 112 ms^50^. Others still have found that modulation of rapid motor responses, on the order of 150 ms, is dependent on task specific visuospatial features^51^. Reward prediction error modulation in saccade latencies were found to be affected on the order of ∼150 ms^42^. A more recent study has found that changes in direction of movement, specifically when a reach target suddenly decreases in expected value, occur on average ∼248 ms after cue^52^. This body of literature detailing the short-latency rapid motor response suggests that the rapid response in relative velocity that we see can be feasibly attributed to tuned sensorimotor reward-based predictions.

To generate these prediction errors, we provided individuals with stochastic reward feedback independent of motor performance. Previous research has focused on either the effect of varying reward levels^53–55^, or the difference between predictable and unpredictable reward^56,57^ on the invigoration of movement. One other work has found a continuous relationship between movement response tempo (i.e. execution time) and reward-contingency belief^58^. Our work in this study details how the reward-vigor relationship in human reaching is continuous with varying degrees of probabilistic expectation as would be predicted based on the utility-maximization theoretic basis^38^. In both explicit contexts where the reward probabilities are stated, and within an implicit context where individuals must learn and experience the differences in expectations, greater reward probabilities were associated with greater average movement vigor.

Beyond the effects of within-trial reward and prediction error, we investigated the relationship between biomechanical effort, subjective choice preference, and movement vigor. Individual willingness to expend effort has been found to be idiosyncratic^59^. We build upon this finding in our current study, showing that individuals differed in their willingness to expend biomechanical effort to accrue monetary reward, and that these differences were demonstrated both in their choices and kinematic behavior during the preceding single-target trials. Specifically, the magnitude of effect of target-direction on choice preference was correlated with the magnitude of the effect on an individual’s movement vigor (Figure 6B).

Aside from reward-mediated invigorative effects of DA, both in movement and deliberation, other research has found the neurotransmitter instrumental in overcoming costs and providing motivation in the face of effortful action^60–64^, which has been shown to influence motor behavior and deliberation ^27,39,65–67^. However, phasic midbrain DAN activity may not necessarily code for upcoming effort value^68^. In monkey (Macaca mulatta) SNc, only a minority 13% of spiking activity in a reaching task was correlated with net utility (rewards minus cost) compared to reward-alone. One hypothesis is that the phasic DA response incorporates effort costs only if reward rate, or discounted value of future action, were meaningfully affected^64^. Other neurotransmitters and pathways may be implicated in utility prediction. Specific serotonin receptor activation in mice was found to increase motivation and vigor alongside an increase in dorsomedial striatum extracellular DA^69^. Thus, the effect of variable effort on the behavioral and dopaminergic response to reward prediction error remains an open question.

Here we use a delta-rule learning model to estimate and quantify the trial-to-trial, subjective value of a given target, motivated by its success in literature modelling prediction error responses in human dopaminergic systems^19,70–73^. We found that movement vigor reflected the learned value, updated from trial to trial^43,58^. A critical assumption in our model is that a value prediction error that integrates reward and effort updates learned value, rather than solely a reward prediction error. Effort too has been shown to influence neural activity in rat ventral striatum and dorsal anterior cingulate cortex (ACCd)^74,75^. Likewise, human ACCd activity was found to reflect the interaction between both expected reward and expected effort^76^. We also modeled effort as additively discounting experienced reward, rather than multiplicatively as in previous work^77^. Additive discounting accounts for the invigorating effect of reward and aligns with the conceptual framework that the brain modulates movement vigor so as to maximize reward rate^38,78^. Our approach is validated in our data where the value prediction error, which integrates learned reward and effort costs, has a significant effect on both the within-trial change in velocity and trial-to-trial changes (Figure 6H,I).

Nonetheless, an alternative approach is one where the prediction error does not incorporate the experienced target effort, which would be modeled as follows:

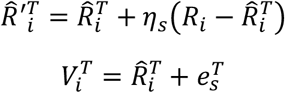

Here, learning is driven by reward prediction alone, and behavior is a result of both learned reward and already known effort. Both models were fit to choice data and compared. The first model, with learning driven from a value prediction error, had greater conditional R^2^, 0.686, compared to the alternative (0.543) when predicting participant choice. However, AICc scores show a preference towards model 2: 2890.61 compared to 2917.76. From our experimental design, the kinematic data would be equally well described with either model, as effort values were constant. Ultimately, the evidence is ambiguous to state conclusively which of the feasible delta-rule models for learning target value best describes the data. Our experiment did not originally set out to test these distinctions, and so additional follow-up with the requisite design considerations would be needed.

The results highlight the differences between an experienced stochasticity and described one^79^, although our first experiment may be better described as a hybrid protocol, as trial-to-trial feedback is still provided to reinforce the previously stated reward expectations. The use of kinematic responses, specifically peak velocity on outgoing and return portions of the movement, are useful tools in understanding not just these differences^80^, but also the similarities between the two environments. The effect of reward prediction error was similar, occurring within the same time frame and showing comparable magnitudes of response. The most apparent distinction was the influence of recent reward history. In our second experiment, the found relationship between prior reward and velocity can be said to have a longer view of the past, integrating incrementally over many trials to arrive in near proximity to the reward average (0.5). The first experiment could be said to elicit more impulsive or rapid response, with the influence of trials further back in time quickly diminishing. This behavior may speak to the difference in environmental uncertainties, where average reward expectation is immediately known to the participant compared to when it must be experienced and learned over time. Previous research has found significant effects of environmental uncertainty in mice and monkey reward learning rates^81^, with greater uncertainty resulting in reducing update rates, which would explain differences in reward history α-coefficients between the two experiments.

## Conclusion

Across two experiments, we demonstrate the sensitivity of movement vigor to dopaminergic correlates of learning and motivation. As the likelihood of reward feedback increased, so too did vigor. Movement vigor was also rapidly updated during an ongoing movement, in proportional response to reward prediction error. Trial-specific learned value, modelled via a delta-rule update formulation, could incorporate both rewards and subjective efforts to predict individual reaching vigor. Lastly, target-agnostic reward history significantly influenced vigor, even when the value expectation was matched. Taken together, our results highlight the link between known short-latency dopaminergic learning signals and the invigoration of movement, not only at the time of cue presentation and movement initiation, but during an ongoing movement immediately after feedback is provided.

## MATERIALS AND METHODS

### Participants

The study consisted of two experiments, each with an independent population (first: n=42, f=16, age=22.3±0.76; second: n=22, f=16, age=23.5±0.9). All individuals were either ambidextrous or right-handed as determined by the Edinburgh Handedness Inventory survey, as well as free of upper extremity injury or self-reported neurological condition. In the first experiment, participants were compensated at a fixed rate of $10 per hour with no difference depending on performance. For the second experiment, the $10/hr rate remained with the potential of an additional $5 depending on performance (average compensation = $14.02±0.13). Participants provided written and informed consent, and procedures were approved of by the University of Colorado Boulder institutional review board.

### Experimental Design

Participants performed a bandit-like task using the KINARM end-point robotic arm (BKIN Technologies, ON, CA) to control a white cursor presented onscreen. From a home circle (r=1cm) at the center of a ring of radius 10 cm, individuals were instructed to make an out-and-back reaching motion to move a cursor (r=0.5cm) a cued target (r=1cm), displayed at 45, 135, 225, or 315 degrees relative to the home circle. Target cues were displayed at a variable time interval uniformly ranging from 800 ms to 1000 ms after trial initiation. On reaching to the cued target, participants need not hit the prompted target exactly. If absolute angular error at the 10 cm radial distance was less than 22°, the target was counted as “hit.” If individuals failed to hit the prompted target, either within the ±22° accuracy constraint or within 4 seconds, a large red “X” appeared onscreen with an accompanying tone (400 Hz, 100 ms duration) indicating a trial failure.

Individuals were instructed that each target had a unique probability (100%, 66%, 33%, or 0%) of providing rewarding feedback at the moment of target hit. Reward feedback consisted of a brief, high-pitched tone (880 Hz, 100 ms duration) being played, the target instantly doubling in size, changing color to yellow, and blinking intermittently (2 blinks, on-off duration at 50 ms). Sample trial progression is shown in figure 1c. Reward probability per target changed between blocks, with a total of 4 blocks each consisting of 180 trials. Trial order and rewards were pseudorandomized such that for each set of 36 trials, each target was cued 9 times, and reward feedback per target was given 9, 6, 3 or 0 times for the reward expectations of 100%, 66%, 33%, and 0% respectively. Individuals were not told the total number of trials to be experienced, only that the total time in the experiment would require approximately 1 hour.

Reward prediction error was defined as the difference between the received feedback and the presented target’s reward expectation, i.e.:

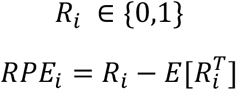

Thus, the four target reward probabilities led to five different reward prediction errors varying in both magnitude and sign. The probabilistic targets (33% and 66%) led to both smaller and larger, positive and negative reward prediction errors. The deterministic targets (0% and 100%) led to 0 RPE, but also allowed us to compare the effects of reward feedback per se, independent of reward prediction error.

In experiment 1, participants were explicitly told which targets were to have what expectation of reward within a block. In experiment 2, the rewards were identical to experiment 1 except that participants were not told of the target reward probabilities but rather had to learn from experience when reaching to the target when prompted. To quantify the degree to which they learned the reward probabilities, the final 36 trials of each block were ‘choice trials’ where two targets were cued rather than a single target. No reward feedback was provided within these choice trials after a target was hit on the reach. Individuals were familiarized with the nature of these choice trials beforehand and were informed that the underlying reward frequencies remained constant within a block. Each unique choice pair (n=6) was presented six times during these choice trial periods. For choice trials, we analyzed the choices as a function of expected value and expected reward. We also measured the response time, the time between target presentation and movement onset. Participants were instructed in the second experiment that additional compensation was contingent on the potential rewards received during these choice trials and thus were incentivized to reach towards targets with previously experienced greater expectation for reward feedback. Total bonus compensation was calculated as the sum of chosen expected reward, divided by maximum potential expected chosen reward (18.66), times five dollars:

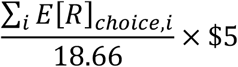

In the first experiment, individuals were familiarized with the reaching task by completing 16 out-and-back reaches, four each to the four different targets. In the second experiment, an additional 8 choice trials were provided as familiarization. Reward feedback was withheld on all familiarization trials.

### Kinematic Metrics

KINARM encoders sampled hand position and velocity (x,y cartesian coordinates) at 1000 Hz. Raw kinematics data were filtered via third-order double pass filter with cutoff of 10 Hz. Radial velocity was computed from numerical differentiation of the radial position relative to the home circle using a second order centered finite difference. The primary metrics on each trial (both single target trials and choice trials) were movement initiation time (reaction time), peak outgoing and return radial velocities, maximum excursion radial distance, and movement duration from onset to target hit. Reaction time, defined as the time between cue presentation and detected movement onset, was calculated by use of the MACC-based onset detection method^82^.

### Trial Exclusion Criteria

For the first experiment, 1.83±0.35% of trials per participant on average were excluded from kinematic analysis either due to failure to complete the trial, failing to reach the outer target ring during the initial outward reach (a “double-peak” movement), or missing the target completely (angular error was ≥22.5°). In the second experiment 7±2.37% of single-target trials per participant on average were excluded from further kinematic analysis. These trials were included, however, for the purposes of calculating reward history and recorded as providing no reward feedback.

### Linear Regression Models

To determine the relationship between velocity, target reward, and reward prediction error, we used generalized Gamma linear mixed models with a log link function to estimate the relative effects of factors of interest and to account for between-participant variability. The Gamma distribution was selected after post-hoc residual analysis of a gaussian-residual assumption revealed significant heteroskedasticity and skewness. The log-link was ultimately selected after model comparisons to different potential link functions (inverse and identity). A similar analysis was used to probe the relationship between reaction time and target reward expectation.

In the first experiment, our regression models for peak velocity and reaction time was as follows:

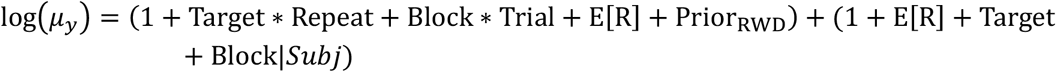

With *µ_y_*representing the mean of the outcome of interest, i.e., velocity or reaction time.

And for the second experiment, an additional interaction term with trial was added:

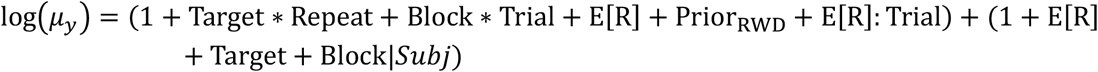

Target is treated as factored variable, and all others as continuous variables. The regression model for return peak velocity was similar, but with added terms for reward reception and prediction error while controlling for outward peak velocity:

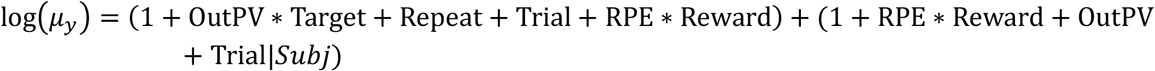

For velocity difference models, a gaussian distribution was found to better approximate the residuals, so a typical LMER with an identity link function was used instead of the generalized Gamma model. All mixed regression models were fitted to restricted maximum likelihood (REML) with the *lme4* and *lmerTest* packages for the *R* language^83,84^, except for when model comparisons were performed with the *performance* R package^85^. P-values for mixed model regression coefficients are derived from Satterthwaite approximations. Hypothesis testing for linear combinations (LHT) of regression coefficients and calculation of asymptotic Chi-squared test statistics was performed with the linearHypothesis function in the *car* R package^86^.

### One-Dimensional SPM Analysis

Continuous time analysis of radial velocity was performed by one-dimensional statistical parametric mapping^87^. Biomechanical curves are well-suited to analysis by this methodology, owing to their smoothness and discrete bounds. We measured the potential effect of reward feedback on different expectation contexts, either deterministic or stochastic, with either single-sample t-tests or 2-factor ANOVAs respectively. This was done to both test for potential significant effects as well as when such effect may occur on instantaneous radial velocity. We focused on the time window from 150 ms prior to hitting the target (i.e. feedback reception) and 300 ms afterward.

Population-level inference on the effect of reward prediction error on within-trial normalized velocity (taken as instantaneous velocity divided by the within-trial peak outgoing radial velocity) was conducted in a manner following previous work^88^. First, regression analyzes were fitted per participant, then this collection of 1-D beta values were submitted to a second-level single-sample t-test.

### Learning Model Design

To model the trial-to-trial update process in experiment 2, we developed a Bayesian hierarchical Rescorla-Wagner learning model. In each of a block’s single-target trials, the presented target’s estimated value was updated based on this rule:

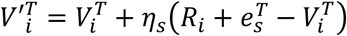

With *T* the presented target at 45, 135, 225, or 315 degrees, *R* the reward feedback [0,1], *e* the effort cost, *s*: participant, *i*: trial, and *η*: learning rate. Participant-specific learning rates (*η_s_*) and effort costs (*e_s_*) were found via posterior maximum likelihood estimation based on choices. The full model (with priors) is characterized as follows:

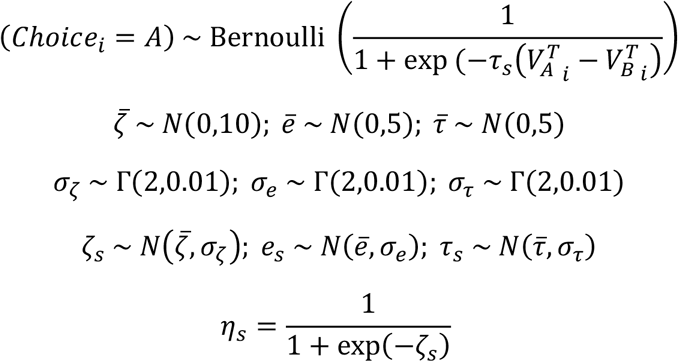

In each choice trial, targets were randomly labelled as either A or B. Value prediction error was defined as the sum of reward feedback and experienced effort minus learned value:

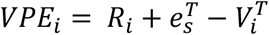

Sampling for Bayesian models was performed by NUTS Hamiltonian MCMC algorithm via use of the *RStan* package^89^.

### Reward History Model

To investigate the effect of prior reward on excursion peak velocity, we applied an exponential moving average to reward reception history^41^. For each single-target trial, average reward was updated with this rule:

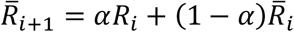

The smoothing factor, *α*, was fit to maximize the likelihood of a reaction time random intercept regression^20^:

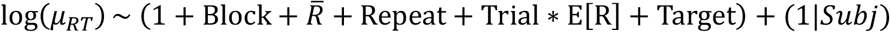

The effects of block number (Block), direction repetition (Repeat), trial number (Trial), reward expectation (E[R]) and factored target direction (Target) were included as control variables. We then used this model-derived reward history term to independently predict outgoing peak velocity in subsequent linear models.

## GRANTS

Work is supported by grants from the National Institutes of Health (1R01NS096083) and the National Science Foundation (CAREER award 1352632) to AAA.

## DISCLOSURES

The authors report no potential conflicts of interest.

## AUTHOR CONTRIBUTIONS

CCK and AAA conceived and designed experiment. CCK performed experiments and analyzed data. CCK and AAA interpreted results of experiments. CCK prepared figures and drafted manuscript. CCK and AAA edited and revised manuscript. CCK and AAA approved final manuscript.

## Notes

### Competing Interest Statement

The authors have declared no competing interest.

### Summary of Updates

Previous research has demonstrated the invigorating effects of reward on movement. Growing evidence suggests this is causally explained by midbrain dopamine transients. Here, we demonstrate that reach vigor tracks canonical variables of learning and motivation across time scales ranging from milliseconds to minutes. Velocity was modulated by reward expectation, reward prediction error and reward rate, key variables that have also been associated with striatal dopaminergic fluctuations. These results point to a potential neural mechanism by which dopamine can influence both decision making and movement control and support the proposition that reward-based invigoration of movement is in part influenced by dopaminergic circuits.

