## Supplemental for "Rapid Dopaminergic Signatures in Movement: Reach Vigor Reflects Reward Prediction Error and Learned Expectation"

### Reaching Vigor Tracks Learned Prediction Error

#### Supplementary Information

##### Post-hoc comparison testing of outgoing peak velocity towards different target directions

*Supplementary Table 1.* Post-hoc multiple comparison testing for differences in excursion peak velocity between the four target directions. Significant adjusted p-values for target velocity comparisons are bolded. Experiment 1.

| PV Comparison | Mean Difference Estimate | Std. Error | z value | Adj. $P_r(>\|z\|)$ |
|---|---|---|---|---|
| $PV_{135} - PV_{45}$ | -0.05206 | 0.005361 | -9.711 | **< 0.001** |
| $PV_{225} - PV_{45}$ | -0.02084 | 0.006761 | -3.083 | **0.01025** |
| $PV_{315} - PV_{45}$ | -0.03804 | 0.006197 | -6.138 | **< 0.001** |
| $PV_{225} - PV_{135}$ | 0.03122 | 0.004717 | 6.618 | **< 0.001** |
| $PV_{315} - PV_{135}$ | 0.014029 | 0.003943 | 3.557 | **0.00212** |
| $PV_{315} - PV_{225}$ | -0.01719 | 0.004047 | -4.248 | **< 0.001** |

*Supplementary Table 2.* Post-hoc multiple comparison testing for differences in outgoing peak velocity between the four target directions. Significant adjusted p-values for target velocity comparisons are bolded. Experiment 2.

| PV Comparison | Mean Difference Estimate | Std. Error | z value | Adj. $P_r(>\|z\|)$ |
|---|---|---|---|---|
| $PV_{135} - PV_{45}$ | -0.06669 | 0.003946 | -16.9 | **< 0.001** |
| $PV_{225} - PV_{45}$ | -0.03109 | 0.003969 | -7.834 | **< 0.001** |
| $PV_{315} - PV_{45}$ | -0.05429 | 0.003958 | -13.716 | **< 0.001** |
| $PV_{225} - PV_{135}$ | 0.035599 | 0.003961 | 8.987 | **< 0.001** |
| $PV_{315} - PV_{135}$ | 0.012404 | 0.003941 | 3.148 | **0.00876** |
| $PV_{315} - PV_{225}$ | -0.0232 | 0.003971 | -5.841 | **< 0.001** |

*Supplementary Table 3.* Post-hoc multiple comparison testing for differences in outgoing peak velocity between the four target directions after application of mixed regression to model the effect of learned value that integrated both reward and effort. Significant adjusted p-values for target velocity comparisons are bolded. Experiment 2.

| PV Comparison | Mean Difference Estimate | Std. Error | z value | Adj. $P_r(>\|z\|)$ |
|---|---|---|---|---|
| $PV_{135} - PV_{45}$ | -0.0663 | 0.01460 | -4.542 | **3.135e-5** |
| $PV_{225} - PV_{45}$ | -0.0306 | 0.0129 | -2.380 | 0.0776 |
| $PV_{315} - PV_{45}$ | -0.0546 | 0.0131 | -4.174 | **1.859e-4** |
| $PV_{225} - PV_{135}$ | 0.0357 | 0.0186 | 1.920 | 0.2128 |
| $PV_{315} - PV_{135}$ | 0.0117 | 0.0150 | 0.783 | 0.8572 |
| $PV_{315} - PV_{225}$ | -0.0240 | 0.0158 | -1.522 | 0.4145 |

#### Single-Target Trial Reaction Time

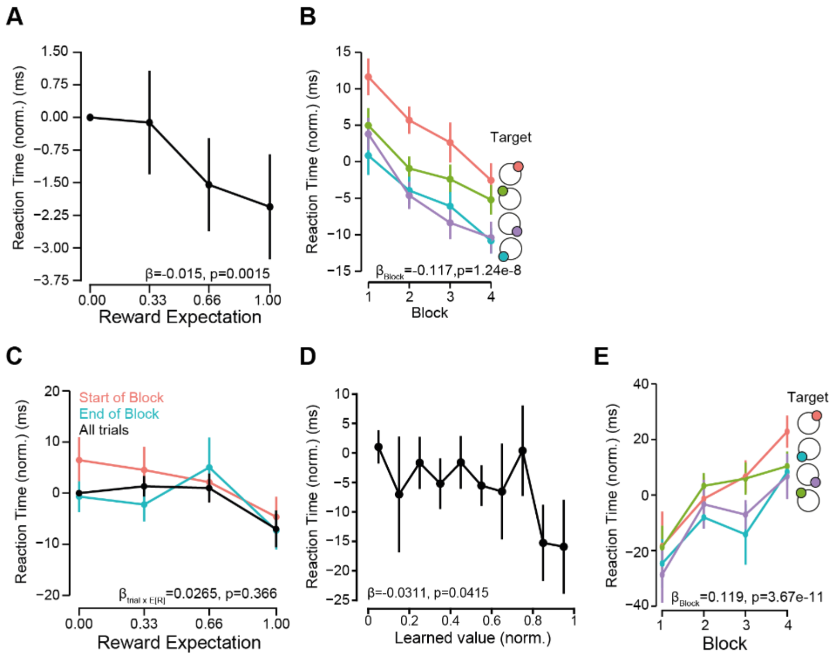


**Figure S1** A) In experiment 1, the time between target cue and movement onset, defined as reaction time, decreased as the cued reward expectation increased. B) Reaction times, over the course of the experiment, decreased, irrespective of target direction. However, reaction times were not stratified in the same manner as was outgoing peak velocities. Evidence suggests that movement initiation times were longer for the top two targets (45° and 135°) compared to the bottom two (225° and 315°). C) In experiment 2, there was no systematic change in relative reaction times for the latent reward expectations over the course of the single-target period. D) Single-target reaction times only marginally decreased as the estimated value for a given cued target increased. E) Unlike in experiment 1, average single-target reaction times increased over the course of the experiment.

#### Experiment 1 Specific Supplementary Results

##### *Control Analysis of Return Movement*

We performed additional control analysis to increase our certainty that it was the online reward feedback, and not other factors, that affected the ongoing movement during the return portion. For this we focused on the two stochastic targets (33% and 66%) that provided a non-zero reward prediction error. The outgoing movements for either target exhibited no difference in peak velocity based on future reward reception (for 33% trials $t_{82}$=0.099, p=0.9212; for 66% trials $t_{82}$=0.0390, p=0.969). For all reaches, the presence of reward feedback also had no effect on the extent of the reach (Gamma GLMM; $\beta_{Reward}$=-0.0485±0.054, p=0.365). Thus, for the same outgoing movement, return movements were faster when a positive, compared to a negative, reward prediction error was experienced. This effect could not be explained by differences in the extent of the reach.

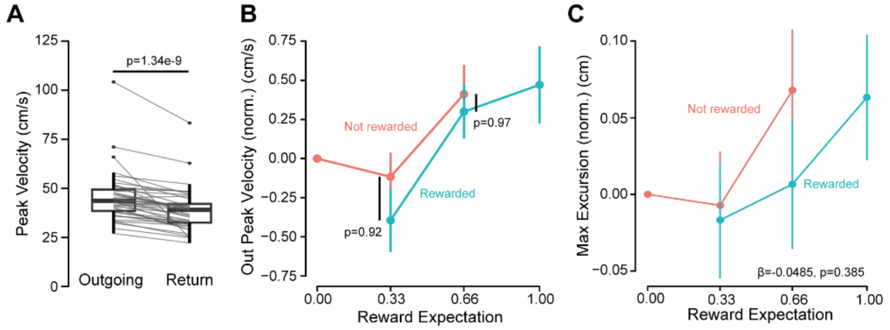


**Figure S2** A) Return peak velocity, on average, was less than the outgoing peak. B) Outgoing peak velocity did not vary with future reward feedback. C) Maximum extent of the reach did not vary with underlying reward expectation, on average, nor did it significantly change after reward feedback was received.

*Effect of Reward on Return Movement*

We were interested in when differences in return velocity due to reward feedback emerged. Difference traces in relative radial position, categorized by RPE condition, suggest an influence on vigor by 300 ms after receiving reward feedback. We also compared instantaneous velocity for rewarding or non-rewarding outcomes, grouped by expectation condition. There appears to be a much greater effect of reward in the return movement within stochastic conditions (E[R] = 66% or 33%) compared to deterministic reward conditions (E[R] = 0% or 100%).

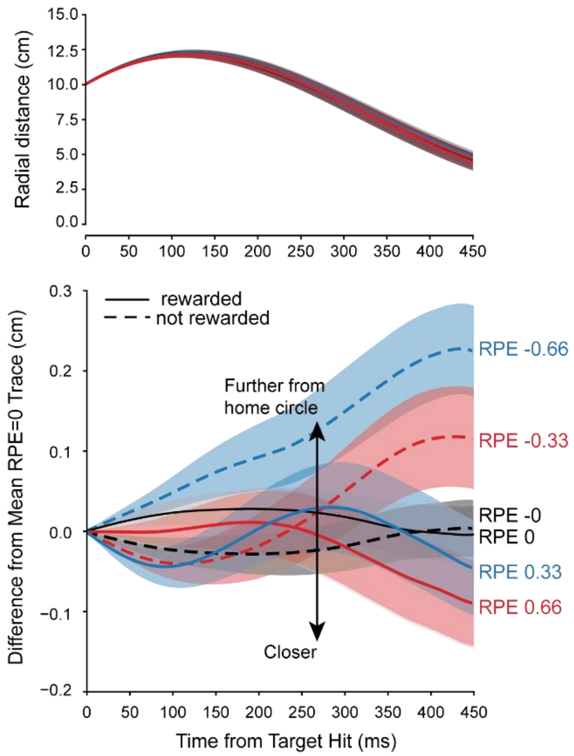


**Figure S3** *Upper* Grand-average radial position traces, aligned to time of target hit, i.e., at 10 cm distance from central home circle. *Lower* Radial position difference traces were aggregated and averaged in a similar manner as in Fig. 2A. For the stochastic reward conditions, we begin to see separation in relative radial position approximately 250 ms after time of target hit. However, for the deterministic rewards, i.e. E[R] = 100% or 0%, no such distinction in radial position after reward feedback is apparent. By 450 ms after time of target hit, individuals are 2mm further away from the home circle after a -0.66 RPE compared to their average for 0% and 100% reward expectation trials. Following a +0.66 RPE, individuals are closer to the home circle by 1mm on average.

#### *Averaged instantaneous radial velocity comparisons*

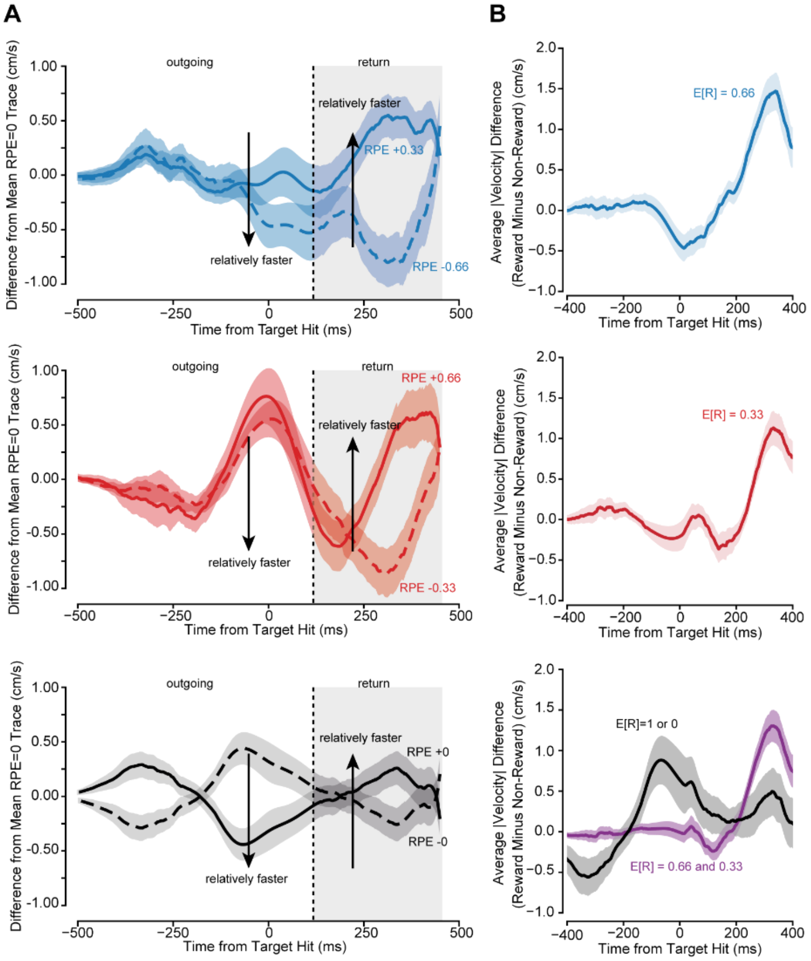


**Figure S4** A) Radial velocity difference trace, aligned to time of target hit, separated by reward expectation and feedback. Colors represent the different expectations and dashed or solid lines represent the absence or presence of reward feedback, respectively. After reward feedback, relative velocities for the same initial expectation differed: more positive RPEs lead to relatively greater return velocities, while negative RPEs led to more negative, i.e. slower, return velocities. Data were subtracted and averaged in a manner similar to figure 2A, but difference was taken to the average of all trials with reward expectation of 1 or 0, rather than only 0. B) Instead of taking the difference to the subject-average for RPE=0 condition, absolute radial velocity data were subtracted from the per-subject average for rewarding and non-rewarding trials for a given reward condition. We see reward feedback produced marked changes in relative velocities approximately 220 ms, on average, after feedback was received. Notably, we do not see a similar magnitude of change for the RPE=0 conditions (bottom).

##### *Return Movement SPM Details*

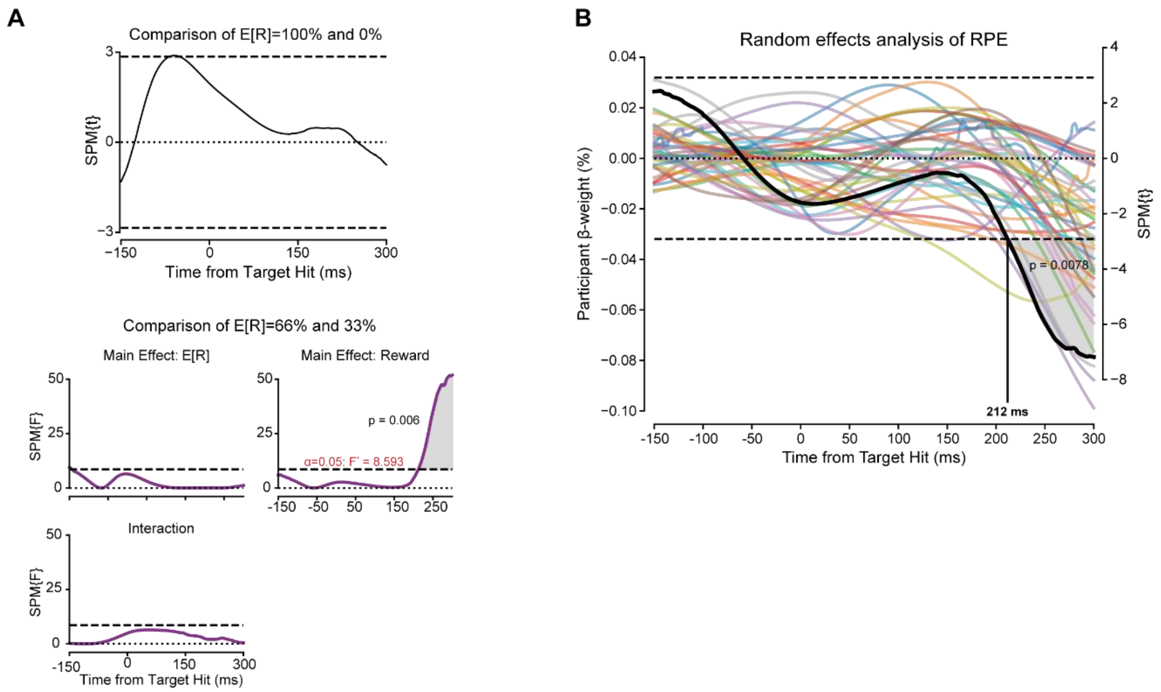


**Figure S5** A) ANOVA SPM results comparing the deterministic and stochastic reward conditions. Single factor analysis reveals no effect of reward feedback in the deterministic case in the time after target hit. For stochastic rewards, 2-factor analysis reveals a significant effect of reward feedback at ~200ms after feedback was received. B) Hierarchical SPM regression analysis reveals a consistent and significant slope effect of RPE on relative instantaneous velocity 212 ms after target hit.

#### Experiment 2 Specific Supplementary Results

##### *Choice Trial Kinematics*

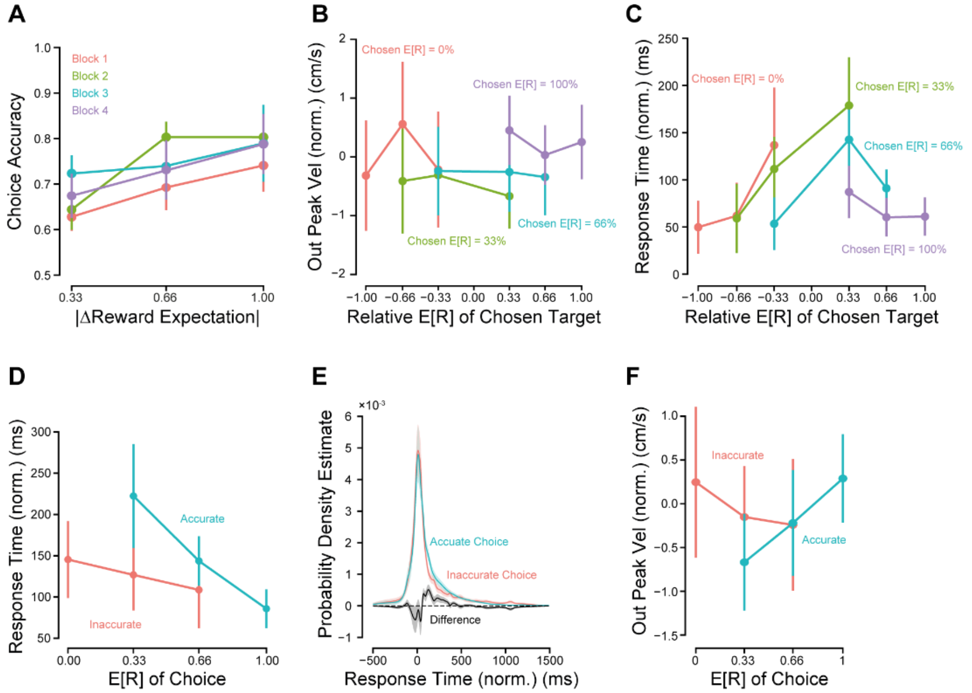


**Figure S6** A) Subject-average choice accuracy, defined as the rate of choosing the more frequently rewarding of the two options presented, increased as the reward difference increased, but did not significantly vary across experimental blocks. B) Outgoing peak velocity during choice trials varied with the expected reward of the choice, but not the relative reward value. C) Response times during choice trials varied with both relative and expected reward amounts. D) Movement onset time during choice trials (response time) decreased as the expected reward of the chosen option decreased. This effect, however, was dependent on choice accuracy, as the slope of response was less for inaccurate choices. E) Distributional analysis of subject choice response times reveals that accurate choices had longer response times compared to inaccurate choices. F) Peak velocity response with respect to underlying reward expectation appeared dependent on relative choice accuracy.

*Outgoing Movement*

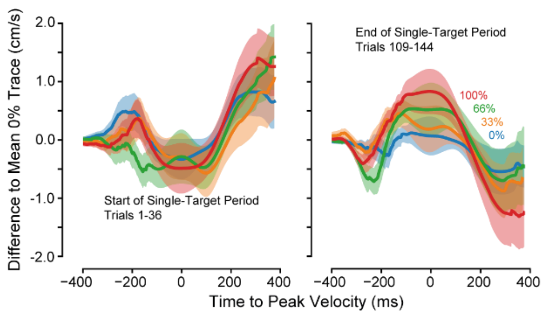


**Figure S7** Velocity difference traces were calculated in the same manner as in figure 2, with the difference taken to the within-subject average velocity trace for the 0% reward condition across the entire block. *Left* In the first 36 trials of the single-target period, there was little if any differentiation between trajectories towards the four different reward expectations. *Right* By the final 36 trials, peak velocity was differentiated between the 4 different reward expectations.

*Return Movement SPM Details*

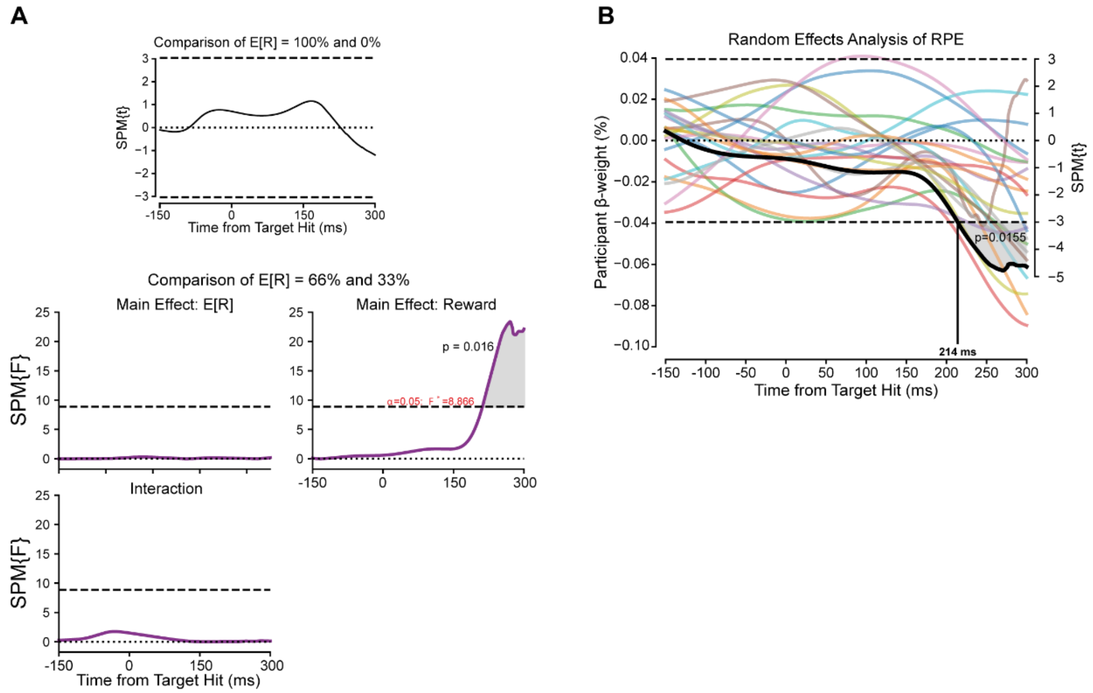


**Figure S8** A) Multiple SPM ANOVA analysis, as in experiment 1, found a significant effect of reward feedback within the stochastic reward trials (E[R] = 33% or 66%). B) Hierarchical SPM regression analysis found a significant effect of reward prediction error on instantaneous radial velocity beginning 214 ms after reward feedback was provided.

*Control Analysis of Return Movement*

As before, we controlled for potential differences in average peak velocity between the outgoing and return portions of the movement, effect of future reward, as well as maximum excursion. We additionally tested whether the effect of RPE on single-target trial return movements were spurious by leveraging the fact that reward feedback is withheld during choice trials and thus providing no prediction error. To test and generate a feasible RPE value, we randomly simulated reward prediction errors for each choice trial depending on participants' selections and underlying reward frequency. After each of the 1000 simulations performed, we regressed this value against the within-trial difference in peak velocities. Following this process, only 85 of the 1000 models produced nonzero regression coefficients with significance ≤0.05. Mean p-value for $\beta_{RPE}$ equaled 0.263 [Wald 95% CI: 0.2357, 0.2903]. In effect, no strong evidence of hypothetical feedback during choice trials was found to predict changes in peak velocity.

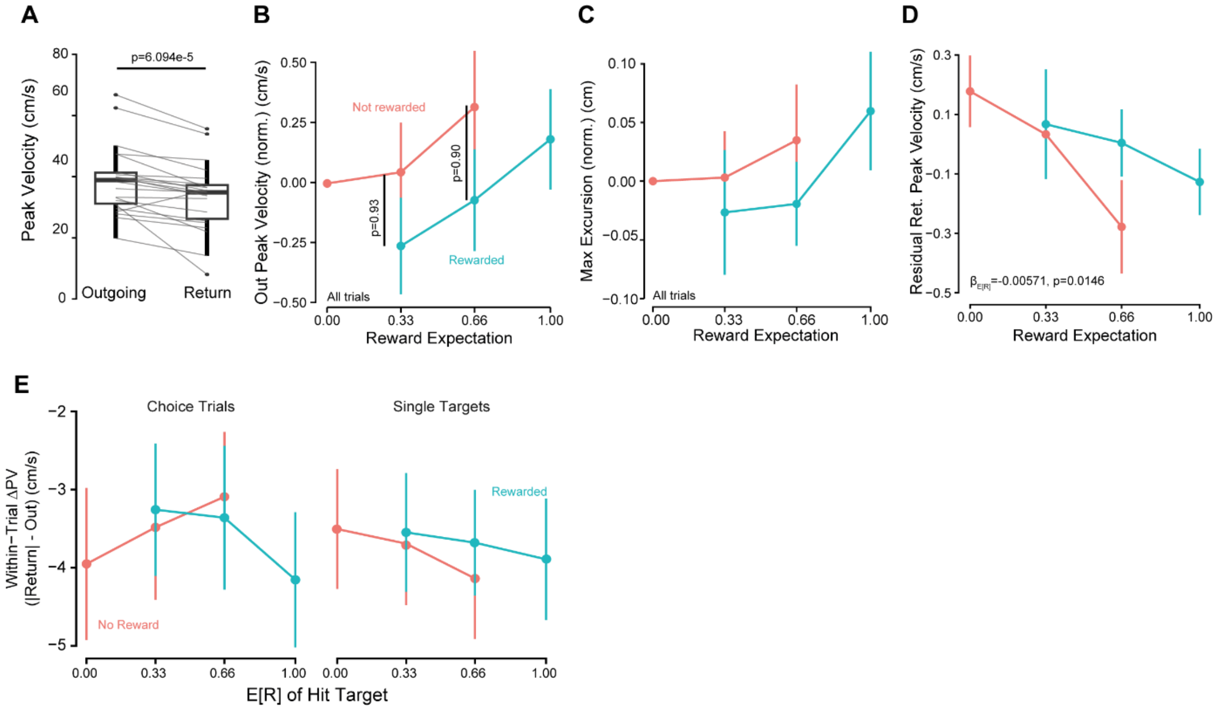


**Figure S9** A) Return peak was, on average, less than the outgoing peak velocity, as in the first experiment. B) Effect of reward feedback or RPE could not be accounted for by differences in outgoing peak velocity. C) Likewise, maximum excursion was indifferent to the reception of reward feedback (β=-3.79e-3, p=0.088). D) Residualized analysis of return peak velocity revealed, as in experiment 1, a significant effect of reward expectation on the return peak. E) Control analysis for the effect of reward feedback on within-trial difference in return peak velocity.

##### *Additional Kinematic Variables Compared against Learned Value Estimate*

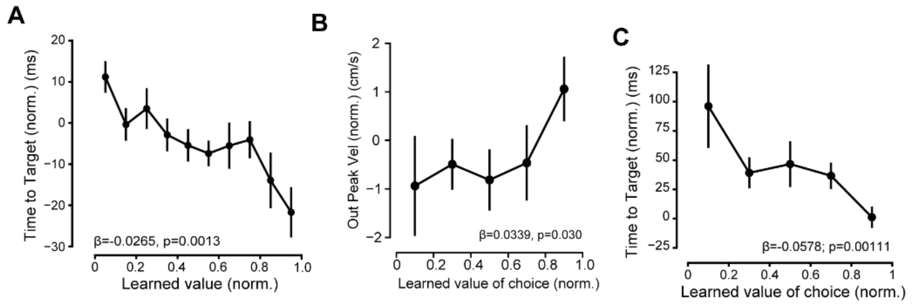


**Figure S10** A) In single target trials, the time to target significantly decreased as the value estimate of a given target option increased. B) Within choice trials, outgoing peak velocity increased with increasing subjective, learned value of the chosen target. C) Similarly, in choice trials, time to target decreased as the chosen value increased.

#### Reward History

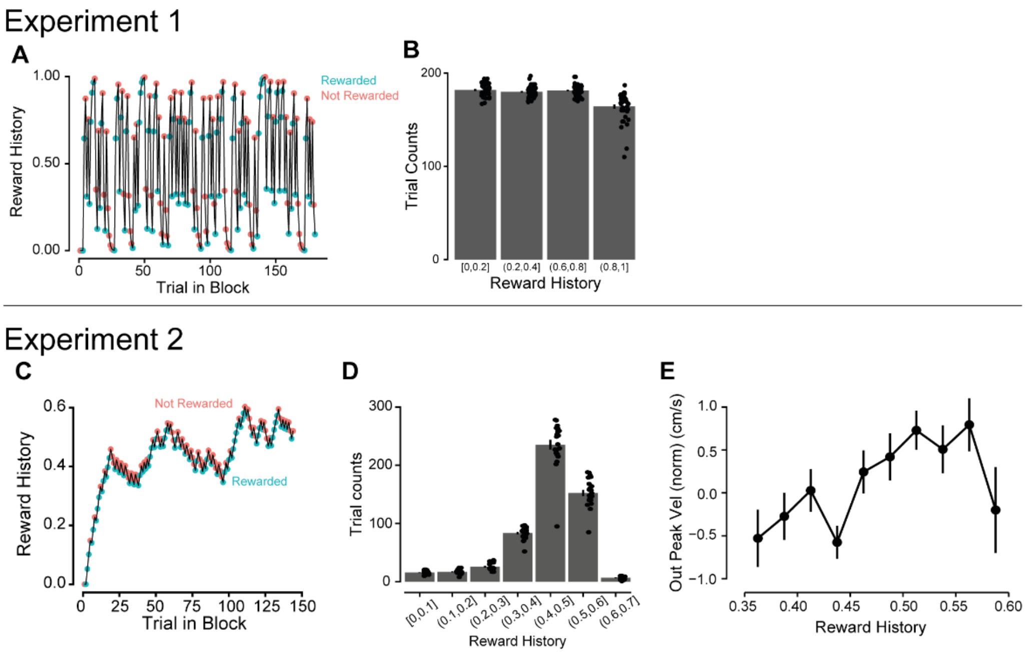


**Figure S11** A) Example reward history values for a single subject and block in the first experiment. B) Averaged trial counts for reward history bins across the subject population in experiment 1. C) Example reward history values for a single subject and block in experiment 2. D) Averaged trial counts for reward history bins in experiment 2. Due to the differences in alpha-coefficient (leaky-integration rate), history values were much less uniform and more concentrated on the expectation mean as compared to experiment 1. E) Highlighting only the final 108 single-target trials, we still see a linear relationship between the reward history value and relative outgoing peak velocity, indicating that this history effect is not driven merely by the initial 36 trials (see *C*).
